# Single-nucleus epigenomic dysregulation unmasks genetic risk-associated neurodegenerative glia states

**DOI:** 10.1101/2025.06.02.657512

**Authors:** Xia Han, Tao Zhang, Chia-Yi Lee, Ashvin Ravi, Salvatore Spina, Alissa Nana Li, Lea T. Grinberg, William W. Seeley, Laura M. Huckins, Towfique Raj, Kristen J. Brennand, Jessica E. Rexach

## Abstract

The accumulation of abnormal tau protein selectively affects distinct brain regions and specific populations of neurons and glial cells in tau-related dementias, such as Alzheimer’s disease (AD), Pick’s disease (PiD), and progressive supranuclear palsy (PSP). Although the three disorders share the feature of tau protein pathology, the regulatory circuitry of non-coding genetic variants underlying risk-associated cell states remains to be elucidated. Using paired single-nucleus profiling of chromatin accessibility and gene expression across AD, PiD, and PSP, we define cell-type-specific cis-regulatory elements (CREs) across six cell types and fifty subclasses. Comparing disease-dynamic CREs across three disorders, we find that glia overrepresent disorder-specific gene regulation related to dynamic cellular response to stress. We show that human genetic variants affecting microglial gene regulation converge into distinct and co-regulated modules affecting specific cellular functions. Moreover, polygenic risk modifiers are maximally co-accessible in disorder-specific glial states, modifying distinct pathways such as sphingomyelin regulation in PiD. Our study informs glial regulators linked to polygenic modifiers of primary tauopathy, introducing modifiable pathways governing resilience for therapeutic consideration.

## Introduction

Neurodegenerative tauopathies are characterized by abnormal tau aggregation in the brain with symptoms of dementia and parkinsonism. Pick’s disease (PiD) and progressive supranuclear palsy (PSP) are primary tauopathies, where tau is the major component of the pathology. Alzheimer’s Disease (AD) is a secondary tauopathy where tau aggregation is regarded to be driven or accelerated by the pathological protein amyloid beta. Tauopathies share a common pathological tau aggregation but vary in symptoms, pathological tau forms ^1^, and genetic architecture ^2^. Neuropathological characteristics distinguish each disorder, including the specific brain regions and cell types most affected by tau pathology and degeneration, and the structure of self-propagating tau fibrils ^1,3^. Genetic risk factors further distinguish each disorder, including several examples of variants with opposing effects on disease risk ^4–8^. Elucidating several diseases in parallel will facilitate our understanding of shared vs. distinct mechanisms linked to disorder-specific genetic and pathological factors across tauopathies.

Although the selective vulnerability of neurons is a major pathological hallmark of disease, glial cells in the brain that maintain and support neuronal functions have been gradually found to be dysregulated in neurodegenerative disease. Recent single-cell genomics studies of AD have revealed signaling pathways disrupted across multiple glial cell types, including inflammation and immune response, lipid signaling, metabolic stress, and DNA damage ^9–16^. Among glia, microglia have gained the greatest attention based on their disproportionate expression of AD-associated risk ^17–21^. Microglial expression of several of these AD risk genes has been associated with beta-amyloid ^19,22–25^. In contrast, glial tau lesions in astrocytes and oligodendrocytes are hallmark features of other tauopathies ^1,26,27^. We recently demonstrated diverse shared and distinct cellular responses to three tau-associated disorders, including an Alzheimer’s-enriched microglia state with high expression of genes associated with AD polygenic risk together with genes known to protect against AD-specific pathology ^19,28^. It is still unclear whether glial subpopulations have diverse roles across tau dementia disorders in disease pathogenesis through distinct mechanisms. Specifically, tau pathology-associated glia in primarily tauopathies remain largely uncharacterized with respect to disease-specific genetics, associated gene regulatory pathways, and downstream functions.

To gain a comprehensive understanding of the epigenomic reactivity and dysregulation of glia subtypes in tauopathies, we conducted an in-depth investigation of paired, single-nucleus profiling of chromatin accessibility and gene expression across AD, PiD, and PSP in three brain regions with distinct vulnerability. By integrating genome-wide association study (GWAS) summary statistics with cell-type-specific chromatin accessibility data obtained across disorders, we enhanced mapping of GWAS variants to functional elements in the human genome and revealed that dynamic chromatin accessibility changes aid in the explanation of disease heritability. Our analysis revealed that the disease heritability of PiD and PSP involved distinct reactive microglial and astrocyte, converging on states exhibiting alterations in sphingomyelin regulation, non-coding circuits relevant to the activation of the lysosome, and shared transcriptional regulators, introducing genetic-risk related glial modifiers associated with resilience to tau pathology.

## Results

To characterize cellular heterogeneity and their epigenomic differences underlying the progression of AD, PiD, and PSP across brain regions, we performed single-nucleus ATAC sequencing (snATAC-seq) on 41 individuals (10 controls, 10 AD, 10 PiD, 11 PSP) across three brain regions: calcarine of the visual cortex (calcarine), insular cortex (insula), and precentral gyrus of the frontal cortex (PreCG) using the 10X Genomics Chromium platform (Fig. 1). This generated a total of 86 samples, comprising 682,667 high-quality individual nuclei after quality control (Supplementary Figs. 1 and 2C and Methods). The overall analysis workflow is shown in Fig. 1. We constructed 8 major clusters after removing 2 undefined groups. The six main cell types were initially annotated based on unambiguous canonical marker expression (Figs. 1B-C, and Supplementary Figs. 2A-2B, and Methods), with 2 excitatory and 2 inhibitory major groups. Marker peaks identified within each cell type showed cell-type-specific TF enrichment. To map chromatin accessibility states to gene expression profiles, enabling cell type validation and deeper insights into cellular regulation, we integrated snATAC-seq with our in-house snRNA-seq using ArchR’s addGeneIntegrationMatrix function ^29^. The snATAC-derived major clusters consistently matched snRNA-derived cell types (Supplementary Fig. 2D), and their cell distribution across brain regions was comparable (Supplementary Fig. 2E). As calcarine samples were limited and showed variability, we focused on the other two regions in the downstream analysis.

**Fig. 1.**
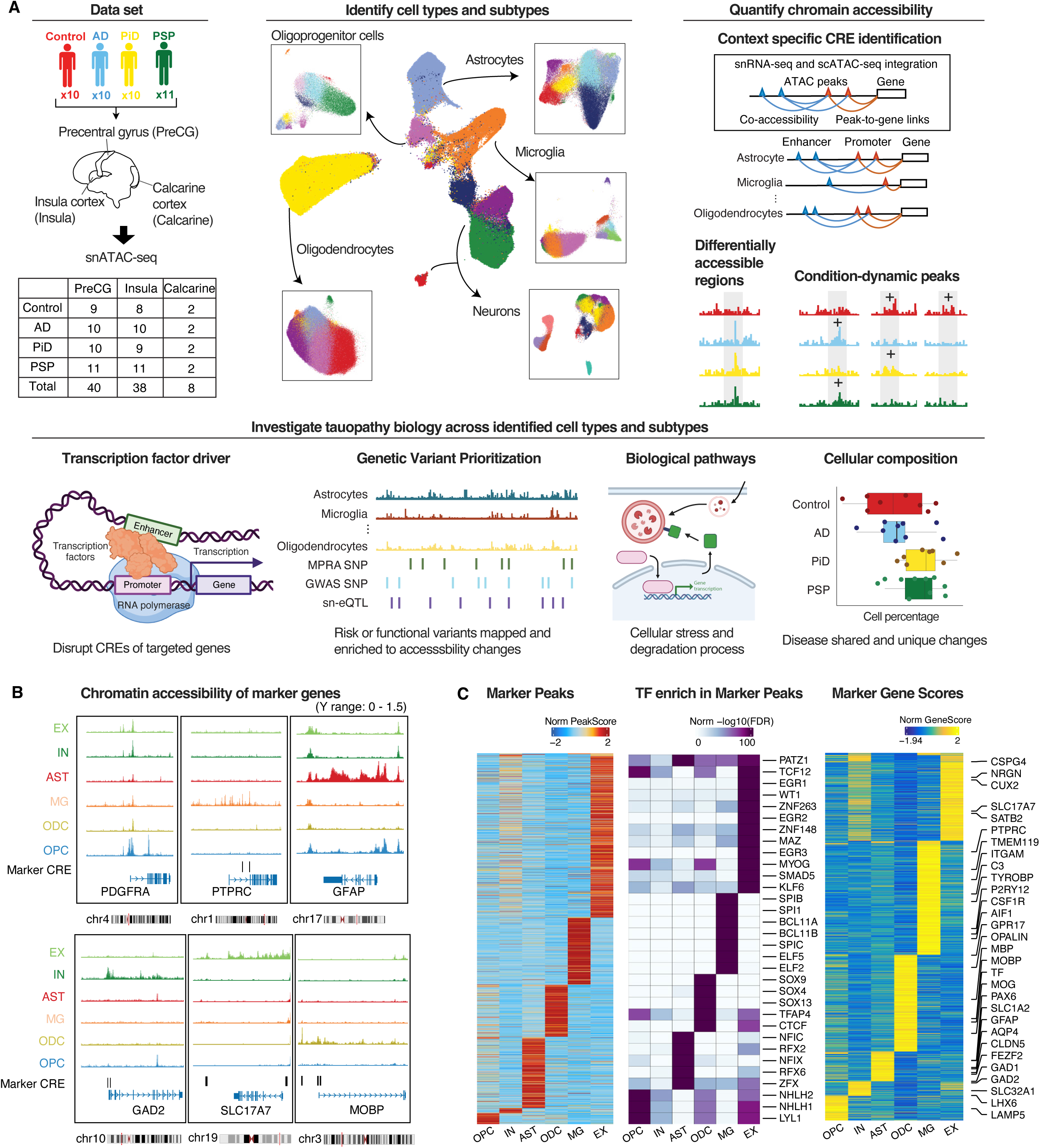
Single-nucleus epigenomic landscape of AD, PiD and PSP across brain regions. (A) Schematic overview of the snATAC-seq analysis workflow. (B) Tracks of chromatin accessibility profiles generated using pseudo-bulk data for each cell type at canonical marker genes. Marker cis-regulatory elements (CREs) of 500 bp are labeled. Visualization and modifications were performed using the UCSC Genome Browser. (C) Heatmaps displaying identified marker peaks (left), marker gene scores (right), and TFs enriched in marker peaks (middle) for each cell type.

Transcription factors are key upstream regulators of gene expression, maintaining cellular viability and functional networks. Changes in TF activity within specific cell types can reflect their regulatory roles and highlight disease drivers. To investigate this, we identified TFs that regulated gene expression and were linked to motif variability in chromatin accessibility (Supplementary Fig. 3A, left panel, and Methods). TFs specific to distinct cell types, such as *SPI1* and *ELF4* in microglia, *NFIC* in astrocytes, *NEUROD2* and *NEUROD6* in neurons, *PBX3*, *SOX4*, and *SOX13* in oligodendrocytes, exhibit consistent patterns of activity, expression, and motif variability (Supplementary Fig. 3A, right panel). Notably, TF variability in PSP glia deviates from other diseases, with glial types clustering naturally within PSP rather than grouping by cell types across diseases. For example, reduced variability in *FOS* and *JUN* was observed across all glial types in PSP (Supplementary Fig. 3B and Methods), suggesting a general PSP-specific TF activation pattern in glia. In contrast, TF variability in excitatory neurons was similar across the three diseases.

## Cell type-specific CREs define cell type identity

Measuring chromatin accessibility on a genome-wide scale enables the definition of potential regulatory elements and illustrates how epigenomic features shape gene expression programs. Using ArchR’s peak calling strategy, we identified a consensus peak set with a total of 924,225 peaks (each 500bp) for all subclusters. To determine which peaks are candidate cis-regulatory elements (CREs) for genes, we combined the peak co-accessibility with gene-peak correlation from snRNA-seq data to infer CREs in cell subclusters split by disease (Methods). We detected 223,710 enhancers and 14,416 promoters. In this case, 25.8% (238,126) of peaks were defined as CREs, improving the interpretation of 7.3% of distal peaks (Fig. 2A and Supplementary Fig. 4A). To validate our candidate enhancers, we collected 11 reference enhancers from public resources, including ENCODE (Encyclopedia of DNA Elements) ^30^, activity-by-contact (ABC) model predictions ^31^, FAMTOM5 (Functional Annotation of Mammalian Genomes 5) ^32,33^, and single-cell studies ^17,34,35^. Based on loci overlapping, we computed the percentages of reference enhancers that were found in our study. As expected, we observed a higher overlapping percentage with known enhancers from single-cell studies ^17,34,35^ than ENCODE bulk data (Supplementary Fig. 4B). In total, 69.4% of our enhancers were supported with the reference enhancers (Fig. 2B and Supplementary Fig. 4B).

**Fig. 2.**
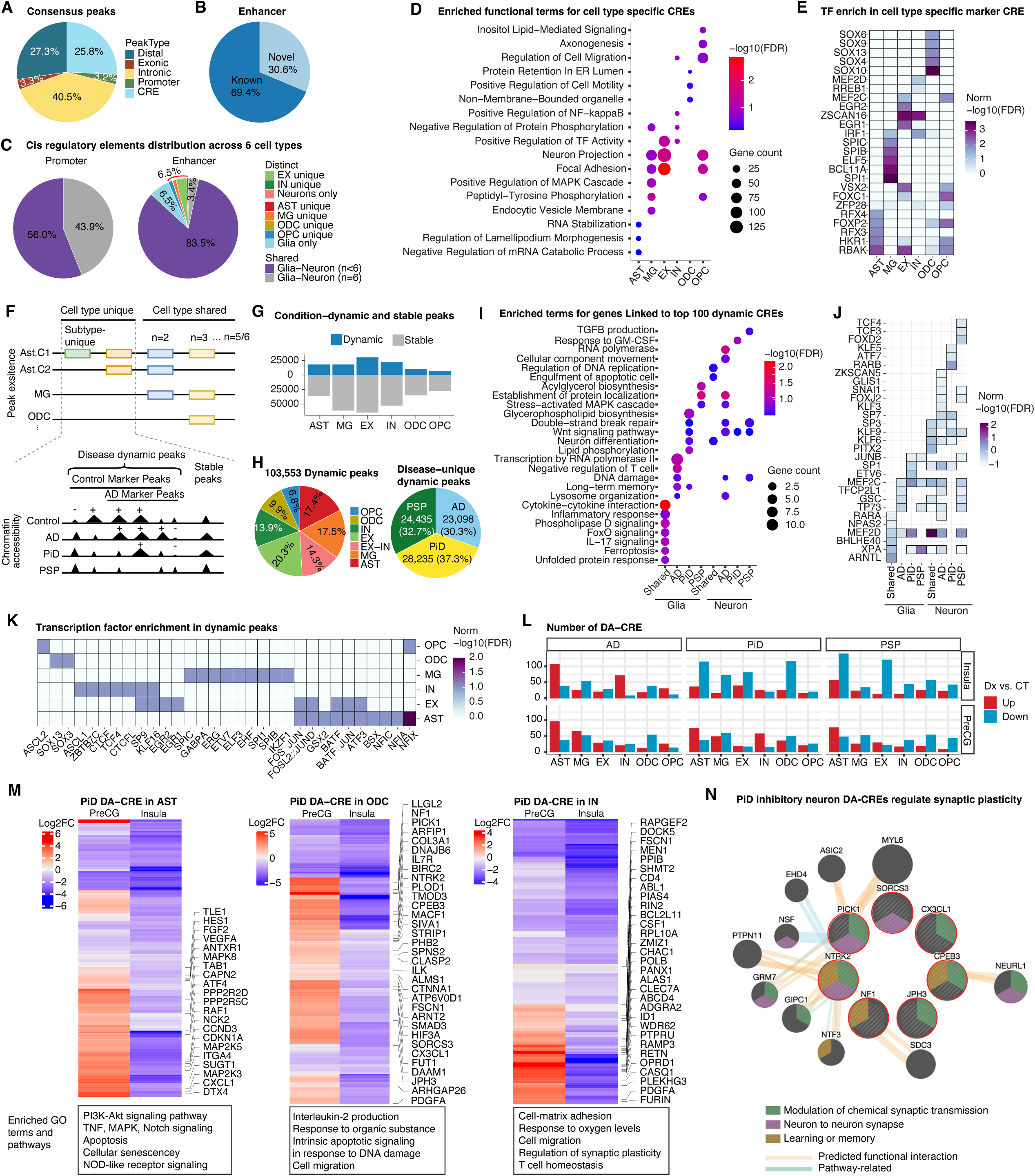
Condition-dynamic and case-control differentially accessible CREs across human brain cell types. (A) Pie chart showing the distribution of consensus peaks across genomic contexts (CRE, promoter, intronic, exonic, or distal regions). (B) Pie chart depicting identified enhancers, categorized as known or novel. (C) Pie chart illustrating the distribution of promoter peaks (left) and enhancer peaks (right) across cell types. (D) Enriched functional GO terms for genes linked to cell type-specific CRE, analyzed using enrichR ^74^. (E) Enrichment of TFs in the top 100 marker CREs uniquely identified in specific cell types, analyzed using MEME ^37^. FDRs were standardized within each cell type group. (F) Schematic diagram illustrating the identification of condition-dynamic peaks for each cell type. (G) Bar plot showing the number of dynamic and stable peaks across cell types. (H) Pie charts display the distribution of dynamic peaks across cell types (left) and across diseases (right). (I) Enriched functional GO terms and KEGG pathways of genes associated with the top 100 dynamic CREs, either unique or shared among the three diseases, in glia or neurons. Functional enrichment was performed using enrichR. (J) TF enrichment for dynamic CREs in the same groups as described in (I), analyzed using MEME. (K) TF enrichment in dynamic peaks per cell type, analyzed using MEME. (L) Bar plot showing the number of differentially accessible CREs per disease across cell types, divided into up-regulated and down-regulated peaks. DA-CREs are identified by P<=1e−3 and |Log2FC| >=1.2. (M) Heatmaps of PiD DA-CREs up-regulated in the PreCG and down-regulated in the Insula, shown for astrocytes (left), oligodendrocytes (middle), and inhibitory neurons (right). Enriched GO terms and KEGG pathways of the genes linked to CREs are displayed at the bottom. (N) Gene network of synaptic plasticity-regulating genes involved in the PiD DA-CREs transition across regions in inhibitory neurons. Genes linked to PiD DA-CREs are highlighted within the red circle. The network is constructed using GeneMANIA ^75^. DA-CREs: differentially accessible cis-regulatory elements.

Sets of persistent epigenetic features establish and maintain specific cell types that share a common genome, where gene promoters are broadly accessible across various cell types, and enhancers are often restricted to particular cell types ^34,36^. We asked how chromatin accessibility in CRE is regulated across cell types. As expected, enhancers were more cell type-specific, while promoters were generally shared among cell types (Fig. 2C). Our findings show that 6.5% of enhancers were unique to only one cell type, and another 6.5% were specific to glia. Functional enrichment of cell type-specific CRE-linked genes revealed the general functions associated with the corresponding cell type identity (Fig. 2D). For example, astrocyte-specific peaks were enriched in regulators of lamellipodia; microglia-specific peaks related to endocytic vesicle membranes; oligodendrocyte-specific peaks were enriched in organelle assembly and cell motility; and OPC-specific peaks enriched in focal adhesion and axonogenesis. Furthermore, excitatory neuron-specific peaks were involved in the regulation of transcription factor activity and neuron projection, while inhibitory neuron-specific peaks unexpectedly related to cell apoptotic process and immune cell activation. Additionally, the cell type-specific CREs were driven by specific transcription factors (TFs) as expected, such as the highest activation of the *SOX10* family in oligodendrocytes and *SPI1* in microglia (Fig. 2E). Importantly, the proportion of annotated enhancers did not differ when comparing our dynamic to stable peaks, and the overall confidence in peak annotation was matched between peak sets.

## Dynamic changes in chromatin accessibility occur across conditions in specific cell types

The epigenomic landscape dynamically controls gene expression in a context-specific manner. Understanding how chromatin accessibility is altered across variable disease-associated contexts elucidates specific pathways and transcriptional drivers of diverse cellular states and functions. First, we identified cell type-specific peaks in the consensus peak set, and we found that 48.7% of peaks were unique in five major cell types (Supplementary Fig. 4C). Next, we identified the dynamic and stable peaks for each subcluster based on whether a peak was a marker for a specific condition (Fig. 2F and Methods). We found that, on average, 16.1% of all cell type-specific peaks were dynamically changed across conditions in different subclusters (Supplementary Fig. 4D). Of the total 103,553 dynamic peaks, approximately half were found in neurons and half in glia, with disease-specific dynamic peaks distributed relatively evenly across all three disorders (Fig. 2H). When analyzed at the cell type level, the average proportion of cell type-specific peaks that were dynamic across conditions increased to 24%, with astrocytes exhibiting the highest percentage of dynamic peaks at 33% (Fig. 2G). This suggests that astrocytes in particular exhibit marked differences in epigenomic dysregulation or epigenomic state transitions when comparing PSP, PiD and AD.

To elucidate common and distinct biological pathways underlying epigenomic dynamics across disorders and cell types, we performed gene ontology analysis on genes linked to dynamic CREs in glia and neurons (Fig. 2I). In glia, disease-shared dynamic CREs were predominantly related to immune regulation, including the inflammatory response and cytokine−cytokine receptor interaction, reflecting a common immune activation in glial cells across conditions. In neurons, disease-shared dynamic CREs were involved in apoptosis and neuron differentiation, and additional disease-unique dynamic peaks regulated stress-related processes, including Wnt signaling and double-strand DNA break repair, which were shared across disorders. Notably, disease-shared and -distinct dynamic CREs and associated biological pathways were more prominent and diverse among glia. For example, disease-shared dynamic CREs in glia regulated inflammation, ER stress and ferroptosis. In contrast, differentially dynamic CREs in PiD and PSP glia regulated lipid metabolism, compared to AD glia which participated more in the regulation of genes involved in T cell activation (Fig. 2I).

To identify the upstream TF drivers of shared and distinct chromatin dynamics, we next performed a TF motif enrichment analysis using MEME ^37^, with disease dynamic peaks as the foreground and stable peaks as the background. For the TF analysis on dynamic CREs in glia and neurons in each disease, we found TFs from the SP and KLF families were enriched in neurons shared across diseases, likely influencing neuronal differentiation and apoptosis (Fig. 2J). In contrast, one of the TFs uniquely enriched in glia is *BHLHE40*, which was known to drive disease-associated microglia response in AD ^38^. To investigate TFs driving dynamic peaks in specific cell types across all diseases, we focused on the top 10 enriched motifs in each cell type and found that disease-associated TFs significantly regulate accessibility dynamics (Fig. 2K). For instance, *PU1* (*Spi1*) and *GABPA* were highly enriched in microglia. *SPI1* was known as a key regulator of microglia activation and AD risk ^39–41^, and *GABPA* was upregulated in PSP and enriched at PSP GWAS-associated disrupted functional variants ^42^. This suggested that while neurons engaged in relatively conserved stress responses across disorders, glial cells displayed more dynamic gene regulation to govern diverse and context-responsive biology.

## DARs indicate a transition in gene activation from middle to high pathology regions

The dynamic accessibility analysis captures broad chromatin remodeling trends across cell types and disease states. To complement this, we performed pairwise comparisons of disease and control samples to identify differentially accessible regions (DARs) at the cell type level, aiming to determine which cell type exhibits the most significant accessibility changes associated with disease (Methods). DARs were identified using the Wilcoxon test in ArchR on the discovery set. To account for the impact of nuclei size on disease outcomes and to ensure the robustness of DAR calling, we performed down sampling 10 times, randomly selecting 30 nuclei per sample for each cell type, and repeated the differential analysis using the Wilcoxon test. In the discovery set, the cell-type DARs have a similar peak type distribution to the consensus peaks (Supplementary Fig. 4E), and 91.6% of the differential enhancers were validated (Supplementary Fig. 4F). Since CREs are more likely to have regulatory potential linked to their associated genes, we focused on differentially accessible CREs (DA-CREs) and observed a higher number of DA- CREs in glia than those found in neurons (Supplementary Fig. 4G, top panel). In addition, astrocytes contain the most significant number of DA-CREs, consistent with our observations of dynamic CREs and further supporting that astrocytes are enriched in disease-associated epigenomic dysregulation. (Supplementary Fig. 4G, bottom panel).

Neurodegenerative diseases exhibit a stepwise progression of pathology from one brain region to another. This progression is accompanied by distinct epigenetic and molecular alterations that vary across specific cell types. Investigating chromatin remodeling across different brain regions, particularly those exhibiting differential tau pathology and neurodegeneration, will provide critical insights into the underlying mechanisms driving the progression of these diseases. In our analysis, both in the original and downsampled datasets, we observed a transition trend of gene deactivation in step with higher pathology when comparing the less affected brain region (PreCG) to more affected brain region (insula), with PiD showing the apparent pattern of down-regulated DARs increasing in astrocytes, inhibitory neurons, and oligodendrocytes (Fig. 2L and Supplementary Fig. 4H). We next examined the functional effects of the genes linked to DA- CREs shared between two regions, which were up-regulated in the PreCG and down-regulated in the insula. In PiD, the deactivated genes in astrocytes were primarily associated with cellular survival and inflammation response, while in oligodendrocytes, they were linked to immune regulation, cell migration, and apoptosis (Fig. 2M). Notably, CRE deactivation in PiD-inhibitory neurons was enriched for genes regulating synaptic plasticity (Figs. 2M-N), suggesting a disruption in neuronal synapse response associated with disease progression. Altogether, the progressive loss of chromatin accessibility in the advanced disease stages underlies the functional impairments observed in glia and neurons and may contribute to the onset of neuroinflammatory response and imbalances in excitatory/inhibitory signaling.

## Disease dynamic peaks improve the explanation of disease genetic variants

More than 90% of the majority of GWAS-identified trait-associated variants are non-coding and likely influence gene regulation through changes in chromatin or cis-regulatory elements (CREs) ^43–46^. While it is known that cell type-specific gene regulatory elements differentially capture heritability factors, it remains unknown whether gene regulatory elements dynamically regulated in disease tissues also differentially capture heritability and if this occurs in a disorder-specific fashion. Therefore, we interrogated whether dynamic peaks in response to disease conditions would capture and interpret heritability differentially than condition-stable cell type-specific peaks. Integrating with GWAS studies of AD, FTD, and PSP, we applied S-LDSC to partition the disease heritability and used two metrics enrichment and standardized effect size (r*) to evaluate the results (Methods). Heritability enrichment is the per-SNP heritability in an annotation divided by the overall per-SNP heritability. r* is the per-SNP heritability correcting for the annotation size and overall heritability.

While individual genetic variants from GWAS studies of AD, FTD, and PSP show similar distribution patterns across both consensus peaks and dynamic peaks (Supplementary Fig. 5A), combined heritability partitioning across different peak sets revealed notable differences between the GWASs. Different cell types captured the greatest amount of heritability of different diseases over disease-dynamic peaks. At the total cell type level, heritability enrichment was most significant across dynamic peaks observed in microglia, inhibitory neurons, and oligodendrocytes in PiD; oligodendrocytes and astrocytes in AD; and neurons for PSP (Fig. 3A). Notably, FTD heritability was most strongly captured by accessible regions whose chromatin accessibility changed in microglia in PiD. This included both gained chromatin accessibility and lost accessibility in a disease context-specific fashion. Peaks that gained chromatin accessibility in PiD microglia specifically showed the greatest overall FTD GWAS heritability score (Fig. 3B). In AD and PSP, significant changes in heritability-associated accessibility occurred in astrocytes. In PSP, peaks that lost accessibility in neurons in disease showed the greatest overall heritability of any cell type, which matches the previous result that GWAS enrichment is lost in subcortical projecting neurons in brain regions where they are selectively depleted, specifically in PSP cases^19^. These findings suggested that disease heritability is influenced by gene regulatory elements that are dynamic in disease-relevant tissue contexts, characterized by the loss of regulatory elements in both neurons and glia during degeneration, and a predominant gain in glia. These alterations differentially affect various cell types in a disease context-specific fashion that further relates to differential cellular vulnerability.

**Fig. 3.**
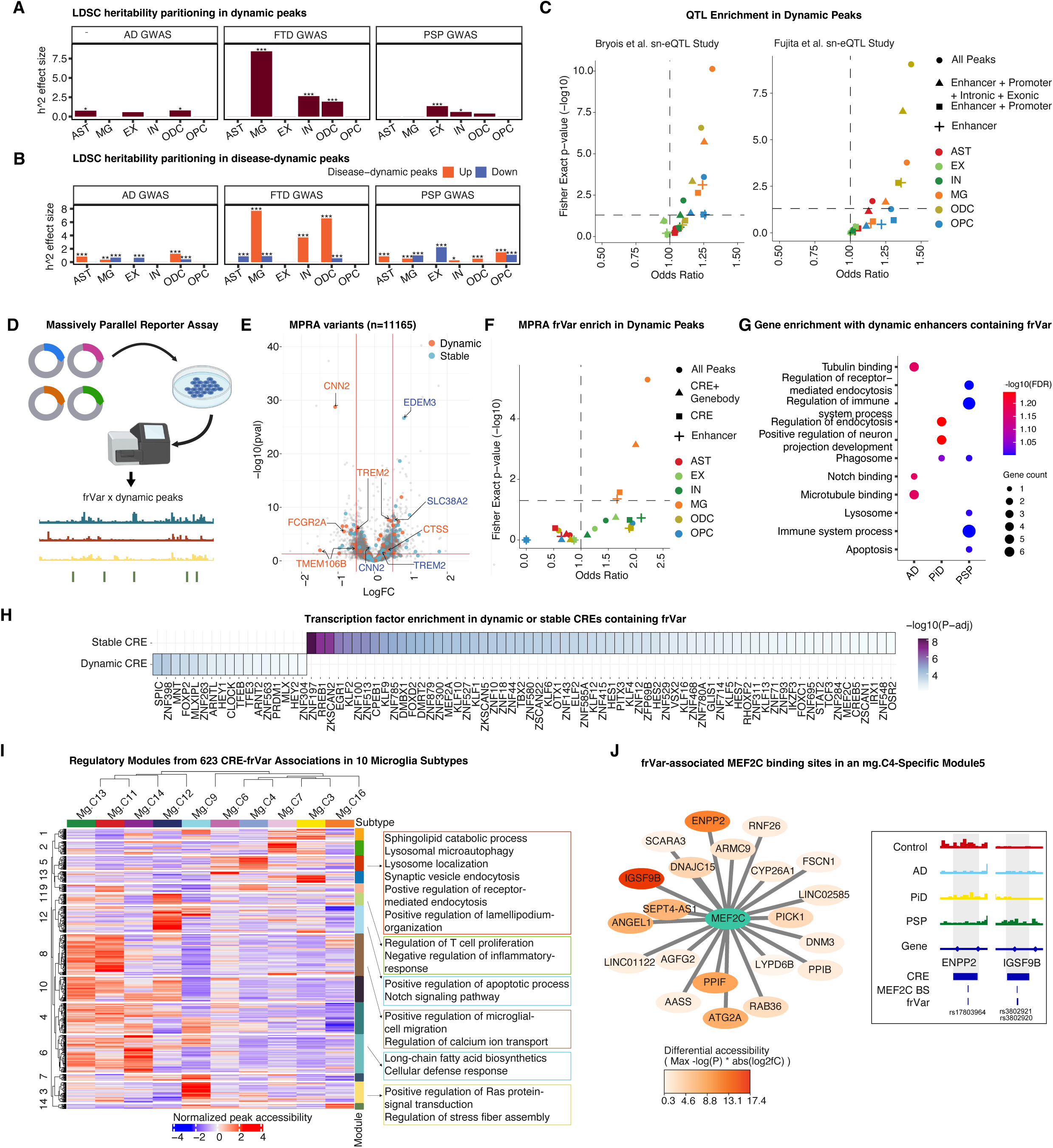
Dynamic accessible regions implicate disease heritability through GWAS, MPRA and sn-eQTL analysis. (A) Partition of disease heritability in dynamic and stable peaks across cell types for AD, PiD, and PSP GWAS, represented by LDSC standardized effect size (r*). (B) Partition of disease heritability in dynamic peaks, stratified by up- and down-regulated peaks for each disease, within each cell type, measured specifically for the corresponding disease. For example, the PSP track depicts LDSC r*for PSP GWAS in PSP up- or down-regulated peaks. (C) Enrichment of sn-eQTLs in dynamic versus stable peaks, tested using Fisher’s exact test. (D) Schematic of the MPRA experiment and integration with dynamic peaks. (E) Volcano plot of MPRA-tested variants, labeled by their overlapping genes’ CREs. (F) Enrichment of MPRA-derived functional regulatory variants (frVars) in dynamic versus stable peaks, tested using Fisher’s exact test. (G) Gene enrichment for dynamic enhancers containing MPRA frVars, analyzed using ShinyGO 0.80. (H) TF enrichment of peaks containing MPRA frVars identified by MEME. (I) Heatmap showing regulatory modules of CREs with frVars across microglial subtypes. Peak accessibility in pseudobulked samples was log2-transformed after depth normalization, and the mean values of subclusters were quantile-normalized. Functional enrichment of CRE-linked genes was analyzed using enrichR. (J) *MEF2C*-target network in mg.C4-specific module5. *MEF2C* binding sites overlapping with frVars are enriched in module5 (bottom). Target CREs in this module are linked to genes enriched for neurodegeneration and endocytosis pathways. CREs linked to target genes are colored by their maximal differential accessibility score, calculated as -log(P-value) x |log2FC| from marker peak calling. LDSC r* with FDR thresholds: *< 0.05; ** <0.005; *** < 0.001.

To test for the reproducibility in independent studies of enrichment for heritability among cell type-specific and disease-dynamic chromatin accessibility peaks, we analyzed snATAC-seq data from the Seattle Alzheimer’s Disease Brain Cell Atlas (SEA-AD) for the middle temporal gyrus (MTG) (Methods) ^47^. We grouped SEA-AD subclasses into six main cell types and identified both condition-dependent peaks and stable peaks within each cell type. We found further AD heritability was significantly enriched only in the disease dynamic chromatin accessibility peaks of microglia (Supplementary Fig. 5B). By examining the SEA-AD data across four conditions (no AD, low, moderate, and high AD pathology), we found that marker peaks in AD samples that were more accessible in low compared to high pathology regions were consistently enriched in AD variants. This supports the conclusion that dynamic peaks related to disease conditions significantly contribute to disease heritability.

## Enrichment of MPRA-validated functional variants within dynamic peaks

Massively parallel reporter assays (MPRAs) are a powerful functional genomics tool enabling high-throughput experimental assessment of the regulatory activity of non-coding DNA elements. To further evaluate the role of dynamic peaks in genetic regulation, we integrated CREs with newly generated MPRA conducted on human microglial-relevant genetic variants in human microglial cell lines, with 11,167 verified variants in total (Fig. 3D, Methods)^48^. We first compared their distribution across dynamic peaks, stable peaks, and the full set of variants, finding that the distributions were comparable (Supplementary Fig. 6A). We then identified 3,325 functional variants (frVars) by p-value < 0.05, with the highest concentration found in microglia enhancers (Supplementary Fig. 6B). Among these, 310 were located in cell type-specific peaks (Fig. 3E). Notably, several neurodegeneration-associated genes, such as *TREM2* and immune-related *CNN2*, were linked to distinct CREs in different cell types in a disease-specific manner. This highlights the importance of context and disease-specific data in identifying candidate functional regulatory elements.

To understand the functional implications of these variants, including their upstream gene regulators and pathways, we conducted transcription factor analysis for dynamic vs. stable peaks containing frVars across all cell types (Fig. 3H). Our analysis revealed distinct patterns of TF enrichment. In dynamic peaks associated with frVars, the TFs identified were predominantly involved in metabolic regulation (*MLXIPL*, *MNT*) and cellular stress responses (*TFEB*, *TFE3*). Additionally, *SPIC*, known for its role in macrophage inflammation and regulation ^49^, was also enriched. These TFs play crucial roles in managing cellular responses and maintaining homeostasis. In contrast, TFs enriched in stable peaks containing frVars spanned a broader range of biological processes, including developmental processes (*FOXD2*, *DMRT2*), cell differentiation (*MEF2A*, *KLF1*), immune responses (*STAT2*, *IKZF3*), and transcriptional regulation (*ZNF197*, *ZKSCAN2*).

We then investigated whether the peaks identified as dynamic in the context of human brain disease are enriched for experimentally validated functional variants, in comparison to peaks that remain stable in disease. We assessed the enrichment of frVars in dynamic versus stable peaks by Fisher’s exact test across different peak types in each cell type. Notably, dynamic peaks in microglia showed significant enrichment of frVars, regardless of the peak type —whether enhancers, CREs, CREs plus gene bodies, or all peaks (Fig. 3F), suggesting that functional variants may modulate gene regulation by perturbing dynamic chromatin regions. To identify which genes were affected in each disease, we performed an enrichment analysis of genes linked to dynamic CREs containing frVars. Both PiD and PSP were enriched for phagosome with PSP- dysregulated genes (*FCGR2A*, *CTSS*) also showing involvement in lysosome and immune regulation. In contrast, AD was significantly enriched in genes related to microtubule binding (Fig. 3G).

To understand the functional implications of these variants in microglia states, including their upstream gene regulators and downstream pathways, we extracted CREs harboring frVar and partitioned them into co-accessible regulatory modules across microglia subtypes using hierarchical clustering based on normalized chromatin accessibility (Methods). These CRE modules exhibited subtype-specific activity and were enriched for distinct biological functions (Fig. 3I). Importantly, module 5, which was specifically activated in mg.C4, was involved in lysosome function, sphingolipid catabolic process, and synaptic vesicle endocytosis (Fig. 3I) and driven by transcription factor *MEF2C* (Fig. 3J). CREs within the mg.C4-specific module harbored *MEF2C* binding sites associated with frVar and were linked to genes enriched in endocytosis and pathways of neurodegeneration-multiple diseases (Fig. 3J). For example, a frVar rs17803964 overlapped a *MEF2C* binding site within a CRE of *ENPP2*, which is involved in the metabolic disturbances in AD ^50,51^. This particular CRE displayed decreased chromatin accessibility in diseases compared to control, while others, including the gene *IGSF9B*, showed increased accessibility. These results highlight coordinated context-dependent gene regulation in microglia, identify high-quality TF drivers based on high-resolution mapping to frVars at nucleotide resolution, and support roles for human genetic variation in more broadly modulating microglial response to stressors in the human diseased brain.

## Enrichment of single-nucleus QTLs in dynamic chromatin

Single-cell resolution quantitative trait loci (QTL) have emerged as a valuable resource to explore the cell type-specific mechanisms that genetic variation influences gene expression in a context-dependent manner. Dynamic chromatin accessible peaks that harbor regulatory variants would support the peaks as functional regulatory elements and reveal various transcriptional activities under specific contexts. To further assess whether disease context-specific dynamic peaks in cross-disorder brain datasets are enriched for regulatory variants based on large datasets sampling common genetic variation and its effects in brain cell gene expression, we leveraged two single-nucleus eQTL datasets from the human brain ^52^ and performed QTL enrichment (Methods). We assessed the enrichment of eQTLs in dynamic versus stable peaks by Fisher’s exact test across different peak types in each cell type. Notably, across different peak categories and cell types, dynamic peaks in microglia and oligodendrocytes consistently exhibited significant eQTL enrichment, particularly within CREs plus gene bodies and across all peaks (Fig. 3C), suggesting that microglia and oligodendrocytes may serve as common targets in tauopathies, where common regulatory variants could modulate gene regulation by perturbing dynamic chromatin regions.

Using CREs harboring eQTLs, we reproducibly identified a mg.C4-specific regulatory module (Supplementary Fig. 6C). Importantly, eQTL module 10, which was specifically activated in mg.C4, was associated with lysosomal membrane function and protein kinase binding. TF analysis over linked enhancers identified nine key regulators, including *EGR1* and *PRDM9*, that drive the activation of module 10, targeting CREs linked to genes enriched in vesicle-mediated transport, lysosomal membrane, and cytoskeleton organization (Supplementary Figs. 6D and S7). For example, the eQTL rs12914843 of *FAN1* (P = 2.26 × 10) overlapped with a CRE of *FAN*1, potentially driving the observed increase in CRE accessibility in PSP and PiD. These findings highlight a potential lysosome-related dysregulation in disease-associated mg.C4. Through the combined eQTL and MPRA-based fVar annotation, we identified a series of candidate TF and target genes poised to regulate microglial lysosomal function and its variation in the human brain. These findings also support roles for human genetic variation in effecting key stress response pathways in glial that are significantly dysregulated in tauopathies.

## Diversity of cellular subclusters across brain regions

Having identified dynamic peaks including functional variants that link to risk genetic factors, we next sought to understand their organization across distinct cell subclusters representing coordinated cellular biological responses to tauopathies. For this, we used the 50 total subclusters that were achieved by combining all cells from a given major cell type across brain regions and disorder for high-resolution cellular subtyping, which we accomplished with negligible batch effect due to our cross-disorder mixed library preparations (Supplementary Fig. 8 and Methods). These included 10 astrocytes, 10 microglia, 12 neurons, 8 oligodendrocytes, and 10 oligodendrocyte precursor cells (OPC) (Figs. 4A-F, Supplementary Figs. 8-9, and Methods).

**Fig. 4.**
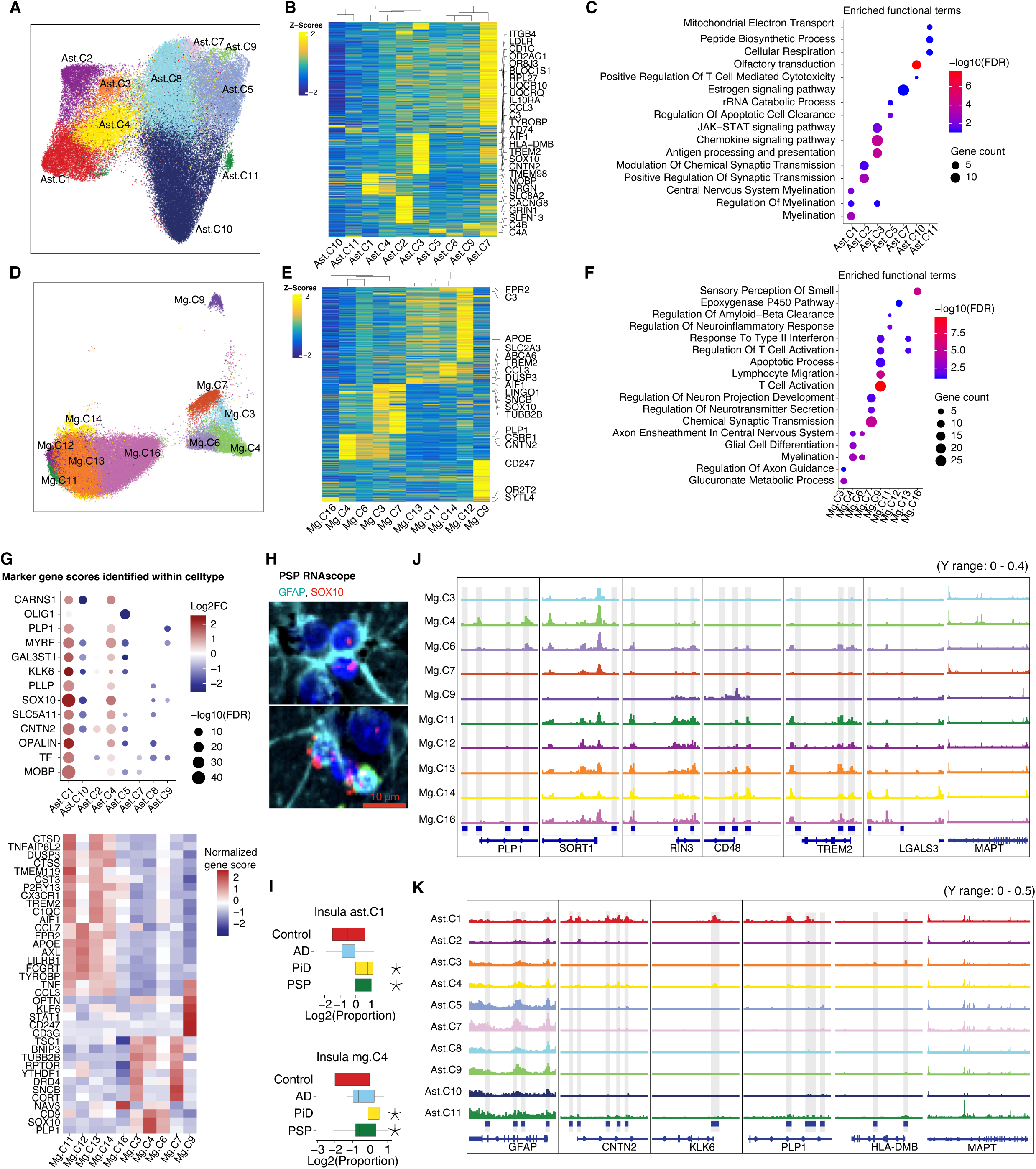
Diversity and heterogeneity of cellular subtypes in human brain of tauopathies. (A-C) UMAP embedding of subclusters of astrocytes (A), with a heatmap of gene score matrix labeled by log2fc > 1 for marker gene scores (B) and enriched functional terms associated with these marker genes (C). ASC, astrocytes. (D-F) Similar analyses for microglia subclusters. MG, microglia. (G) Gene signatures of astrocyte subclusters (top) and microglia subclusters (bottom), defined by marker gene scores compared within each respective cell type. (H) RNAscope ISH for *SOX10* combined with GFAP IHC in human insular tissue from a PSP patient. Representative images show co-localization of *SOX10* mRNA and GFAP in a subset of astrocytes (Scale bar: 10 µm). (I) Boxplots displaying the relative abundance of insula ast.C1 (top) and mg.C4 (bottom) across conditions. Changes in cell composition among disease and control groups are modeled using linear regression computed by Limma, adjusted for age and post-mortem interval. *P<0.05. (J) Example tracks showing marker peaks for genes *PLP1*, *SOT1l* in mg.C4; *RIN3* and *TREM2* in mg.C13 and mg.C6; CD48 in mg.C9 and mg.C6; and *LGALS3* in mg.C7 and mg.C14. (K) Example tracks showing marker peaks for genes *CNTN2*, *KLK6*, and *PLP1* in ast.C1; *HLA- DMB* in ast.C3; and GFAP in multiple astrocyte subclusters.

Consistent with previous studies ^10,12,15,53–55^, we reproduced astrocyte subclusters associated with homeostasis, reactive states (ast.C5, ast.C7, ast.C8, ast.C9), synaptic transmission (ast.C2), antigen presentation (ast.C3), inflammation (ast.C10), and metabolism (ast.C11), as well as microglia subclusters relate to homeostatic (mg.C16, mg.C14); inflammatory (mg.C13, mg.C11); disease-associated microglia (DAM) (mg.C12), synaptic transmission (mg.C4, mg.C7) and T cell activation (mg.C9) (Methods). Additionally, we identified two myelin-related astrocyte subclusters (ast.C1, ast.C4) and microglia subclusters (mg.C4, mg.C6) expressing *PLP1* and *SOX10* (Fig. 4G, Supplementary Figs. 9B-C), extending the findings of prior studies. Marker gene expression in ast.C1 was validated by RNAscope in situ hybridization for *SOX10* and *PLP1* combined with immunohistochemical staining for GFAP protein, in FFPE human insular tissue (Fig. 4H and Supplementary Fig. 10). The tau-encoding gene *MAPT* was upregulated in mg.C4 (Fig. 4J and Supplementary Fig. 11A, left panel) and differentially expressed in PSP in insula in ast.C1 (Supplementary Fig. 11A, right panel), suggesting that these two subclusters may be associated with tau dysregulation.

Chromatin tracks revealed that marker peaks of marker genes were consistently up-regulated in distinct subclusters, such as *GFAP* in reactive astrocytes; *HLA-DMB* in ast.C3 (Fig. 4K); *RIN3* in mg.C13, *CD48* in mg.C9, *TREM2* in mg.C13 and mg.C11, and *LGALS3* in mg.C7 and mg.C14 (Fig. 4J). Both ast.C1 and ast.C4 were myelination-related and oligodendrocyte-like, with ast.C1 displaying prominent marker peaks for *KLK6* and *PLP1* (Fig. 4K). Compared to other astrocyte subclusters, ast.C1 highly expressed oligodendrocyte markers, including *MOBP*, *TF*, and *OPALIN*, as well as disease signature genes found in oligodendrocytes, such as *CNTN2*, *SLC5A11* (Fig. 4G). Similarly, *PLP1,* a X-chromosome gene that participates in age-related resilience ^56^, was most highly upregulated in mg.C4, while mg.C6 exhibited greater chromatin accessibility around *RIN3*, *CD48,* and *TREM2* (Fig. 4J).

For neurons, we identified four inhibitory neurons and eight excitatory neurons (Supplementary Fig. 9B). The four inhibitory neurons consist of four *SLC32A1*^+^ *GAD1*^+^ cells. Among them, three expressed additional markers: *PVALB*^+^ (neu.C7), *SST*^+^ (neu.C8) and *ADARB2*+ *VIP*+ (neu.C6). The four excitatory neuron subclusters expressed layer-specific markers: neu.C5 and neu.C11 were *FEZF2*^+^, neu.C14 was *RORB*^+^, and neu.C13 was *CUX2*^+^.

To contextualize our findings with prior published work, we compared these clusters with previously defined microglia states at the RNA level within this same dataset ^19^ to identify overlapping and similar states (Supplementary Fig.11B). ATAC data provided higher-resolution partitioning of cells compared to RNA data in several instances, notably with mg.C4 and mg.C6, two distinct ATAC-defined clusters that both overlap with the RNA-defined insula-microglia-3 cluster, resembling myelin-eating microglia observed in multiple sclerosis (MS) datasets ^55^, all of which share *PLP1* as a marker gene. While both mg.C4 and mg.C6 share markers of myelination, they differ predominantly in the expression of lipid transporters (*ABCA2*, *ABCA8*, *SCAP*) and lipid metabolism genes (*HMGCS2*, *LPGAT1*), and genes associated with buffering metabolic stress (*SGK2*, *TSC1*). Comparing mg.C4 with the microglia subtypes described in ^55^, we found significant marker overlap with a myelin-phagocytosing microglia population observed in MS (Supplementary Fig. 12A and Methods). However, our cluster demonstrated increased chromatin accessibility at myelin regulatory and processing genes (Figs. 4G and 4J), with co-enrichment at their promoters of *SOX10*, suggesting active regulation in mg.C4. Furthermore, upon re-clustering our in-house snRNA-seq microglial cells into 10 clusters, we identified a microglial subtype expressing both *SOX10* and *PLP1*. This subtype shows a strong alignment with the ATAC subcluster mg.C4 (Supplementary Figs. 12B-C), further supporting that the mg.C4 is putative myelination-related microglia.

## Variable disease-enriched glial states by disorder

We next explored how cellular composition changes in disease within each brain region. Changes in cell proportions by cell type, disease, and brain region were summarized in Supplementary Table 1 and Supplementary Figs. 13-14, with notable examples shown in Fig. 4I. In the insula, we observed a proportional increase in total microglia in AD and in astrocytes in PiD (Supplementary Fig. 13B). At the subcluster level, we identified disease-specific and common subcluster composition changes in a region-specific manner (Supplementary Fig. 14).

We noticed prominent changes in PiD among oligodendrocytes and myelin-related subtypes. PiD exhibited the most pronounced disease-specific changes in subtype composition in inflammatory oligodendrocyte subclusters within the insula, characterized by a decrease in odc.C7 and odc.C8 and an increase in odc.C9. Notably, alterations in myelin-related *PLP1+* subclusters were shared by diseases in the insula but involved different glial types depending on the disorder. Both ast.C1 and mg.C4 were significantly increased in PiD and PSP (Fig. 4I), while mg.C6 was elevated in both PiD and AD. At the same time, Ast.C1, mg.C4, and mg.C6 express their expected markers of astrocytes and microglia, such as *GFAP* and *C3* (Supplementary Fig. 9B). The increase in ast.C1 and mg.C4 in PSP and PiD was also supported by bootstrapped analysis (Supplementary Fig. 11C and Methods). These combined observations of increased accessibility of genes associated with myelination, including *PLP1,* which has recently been shown to be neuroprotective ^56^, in microglia and astrocytes, suggest a shared feature of primary and secondary tauopathies. These findings showed that the insula accumulated pathological signals with the myelin-related microglia and astrocytes increasing in PiD and PSP.

## TF drivers of disease-reactive microglia and astrocytes in PiD and PSP

Importantly, microglia containing *PLP1* mRNA have been reproducibly described in microglia ^55^, and have been shown experimentally to occur secondary to myelin phagocytosis. Our findings suggested that *PLP1* chromatin accessibility varies within disease-associated reactive glia. We aimed to further investigate the gene regulatory mechanisms driving *PLP1* and other myelin-related genes in tauopathies and explore how cell stress and injury pathways interact with tau pathology.

We performed TF analysis and found the master regulator of myelination genes, *SOX10,* was significantly enriched in both mg.C4 and ast.C1 (Supplementary Table 2). *SOX10*, a transcription factor typically associated with oligodendrocytes but also with known roles in neuroprotection, is among the top markers distinguishing ast.C1 and mg.C4 from other astrocytes and microglia. It is also expressed in oligodendrocytes, as expected, and is notable at higher levels (Supplementary Fig. 9B). To identify TF targets by mediating binding events in accessible chromatin, we predicted TF binding sites on the marker peaks of subclusters (Methods). Importantly, while 34.4% of *SOX10* targets overlapped across oligodendrocytes, astrocytes, and microglia, including the myelin-associated genes *PLLP*, several targets were uniquely prominent among *SOX10* targets in the non-oligo cell types, including *PDGFA* and *RBPJ*, supporting shared and distinct effects of *SOX10* by cell type (Supplementary Table 3). In mg.C4, *SOX10* shared targets (such as *CHRAC1*, *API5*) with another enriched TF, *FOXP2*, which is associated with language development and linked to neuropsychiatric disorders, and has been shown to exhibit human-specific expression in microglia ^57^. In contrast, in ast.C1, *SOX10* shared targets (such as *BEST1, CDH5*) with a different TF, *NFIX*, which is an astrocyte-specific TF with roles in astrocyte development and maturation levels (Supplementary Table 4).

Interestingly, *SOX10* target genes in mg.C4 and ast.C1 both enriched in process of cell population proliferation, developmental growth and lipid biosynthetic process (Supplementary Table 5). This suggests that these two clusters, driven by *SOX10,* may play a role in promoting cellular growth and lipid metabolism to maintain brain function. Importantly ast.C1 shows the highest gene activity and expression of *MAPT* compared to all other astrocytes in PSP (Supplementary Fig. 11A and Supplementary Table 6). We next identified TF target genes in each disorder by calculating gene-TF linkage scores within ast.C1 (Methods). We found that *SOX10* was differentially activated in the PSP insula, targeting a SNARE protein, *STX4,* in all disease conditions (Supplementary Fig. 15A). In PSP, the predominant tau species retains exon 10, known as the 4R isoform. SNARE genes mediating lysosomal membrane fusion affect tau propagation in iPSC models of 4R tau ^58^. Therefore, in ast.C1, *SOX10* participates in regulating both myelin genes and pathways previously implicated in the toxicity of 4R tau.

Overall, targets and pathways of *SOX10* across and within each cell type support a generalized role in regulating changes in lipid metabolism in the context of disease, and generally beneficial roles that would support cell survival in the context of stress. This aligns with known neuroprotective roles for *SOX10* in neurons, as demonstrated by *SOX10* overexpression in both CRISPR screens ^59^ and in vivo studies of neurodevelopment ^60,61^. Consistently, epigenomic erosion analysis using single-nucleus methyl-3C sequencing data revealed significant gains and losses of heterochromatin across multiple astrocyte states in PSP, where ast.C1 remained unaffected (Supplementary Fig. 15C, Methods). The maintenance of epigenomic stability in ast.C1 may protect them from depletion as a resilient state.

## Genetic heritability enriched in disorder-divergent cell states in PSP and PiD

Given emerging evidence that polygenic trait expression is most pronounced in disease-specific biological states related to causal pathways, we next employed heritability partitioning to assess relationships between gene regulation, enhancer co-accessibility, disease risk loci, and downstream pathways using diverse disease contexts. Therefore, we performed LDSC as previously described, applied to cell subclusters representing distinct states.

We applied LDSC analysis at the sub-cell-type level for all 50 clusters. Using FTD GWAS as input, we found two clusters (mg.C4, opc.C4) across all cell types were enriched for FTD heritability using LDSC enrichment (Fig. 5D and Supplementary Fig. 15D). Using PSP GWAS as input, LDSC enrichment in clusters did not pass the same P-value significance threshold, while the trend matches to the LDSC r*. The positive and significant r* implied that the PSP heritability enrichment effects were captured by five neuron subclusters (neu.C5, neu.C8, neu.C9, neu.C12, neu.C13), as previously seen at the major cell type level, and two astrocyte clusters (ast.C1, ast.C10) (Supplementary Fig. 15F). Importantly, the excitatory neurons demonstrated an overall loss of chromatin accessibility at PSP GWAS loci, while astrocytes in contrast exhibited a gain in accessibility at these loci (Fig. 3B).

**Fig. 5.**
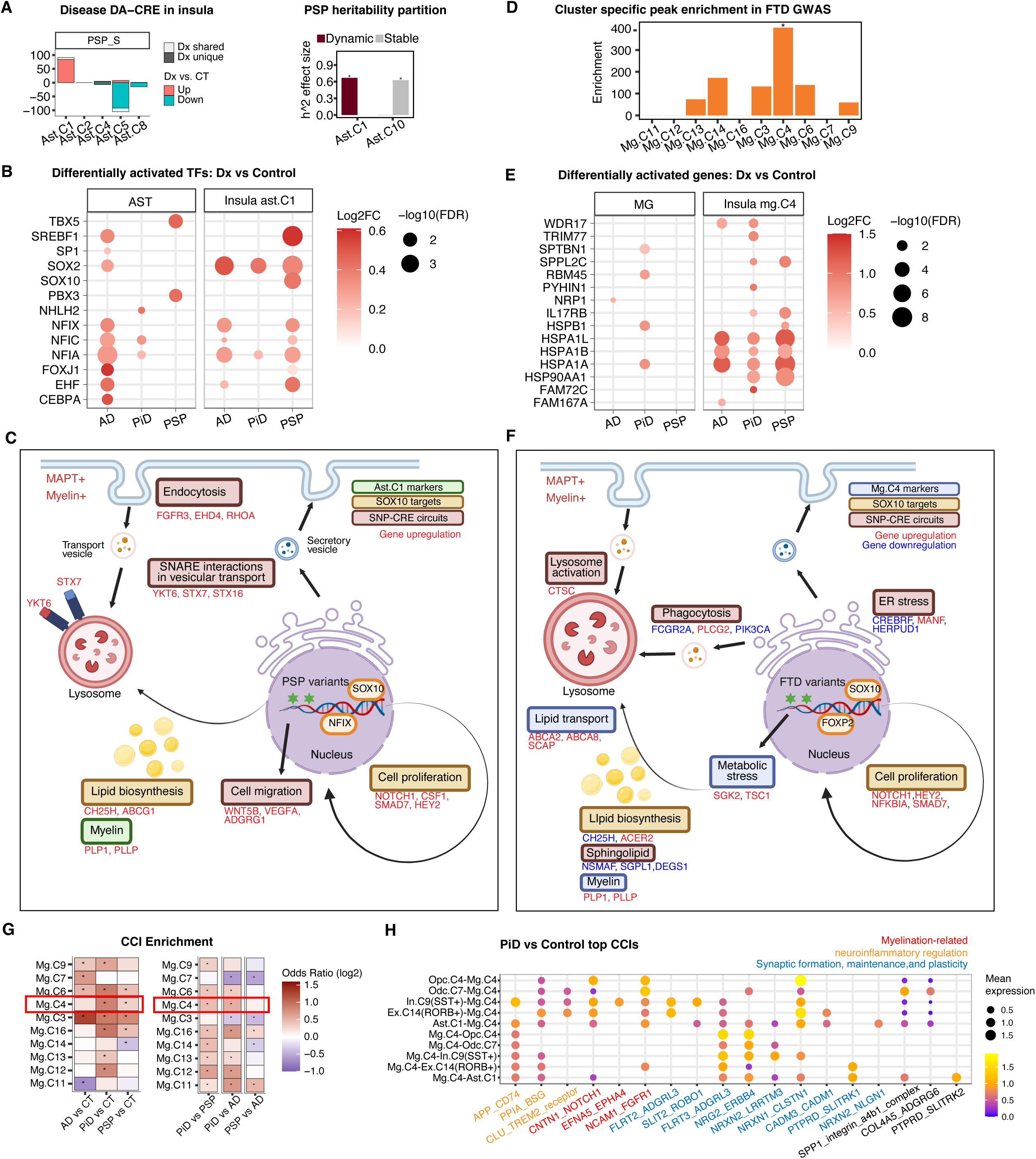
Integrated analysis of accessibility changes, gene activity, GWAS heritability partition, epigenomic stability, and cell-cell interactions identifies disease-associated glia subtypes. (A) PSP-associated chromatin accessibility changes in astrocytes. Bar plots (left) show the number of differentially accessible CREs in PSP across astrocyte subclusters, categorized by up- and down-regulation. The right panel shows partitioned disease heritability of dynamic peaks in ast.C1 and ast.C10 (right), displaying LDSC standardized effect size (r*). FDR *< 0.05; ** <0.005; *** < 0.001. (B) Differentially activated TFs in astrocytes and ast.C1. (C) Schematic of molecular changes and dysregulated pathways driven by PSP GWAS risk variants in PSP ast.C1. Myelin-related astrocytes proliferate in tauopathy, activating SNARE-mediated vesicle trafficking and lysosomal pathway to mitigate lipid stress induced by PSP risk variants. (D) Disease heritability partition in subcluster-specific peaks for FTD GWAS across microglia subtypes. FDR *< 0.05; ** <0.005; *** < 0.001. (E) Differentially activated genes in microglia and mg.C4. (F) Schematic of molecular changes and dysregulated pathways driven by FTD GWAS risk variants in PiD mg.C4. Myelin-related microglia proliferate in tauopathy, activating lysosome- phagocytosis pathways to counteract ER and metabolic stress induced by FTD risk variants. Genes with upregulation are shown in red font, and those with downregulation in blue, based on differential gene scores or genes linked to differentially accessible CREs. (G) Number of cell-cell interactions (CCIs) involving subclusters that are overrepresented in pairwise comparison. (*) FDR < 0.05; chi-square test. (H) Top ligand-receptor pairs mediating interactions between mg.C4 and the represented subclusters in PiD. Values represent the mean expression of ligand-receptor pairs in corresponding cellular pairs.

Our goal was to identify cell states with the most pronounced aggregated co-regulatory structures affecting potential causal SNPs. We aimed to map their key regulators, targets, and downstream pathways and compare these features across disorders. At the sub-cell-type level, we found DA- CREs concentrated in glial subclusters across all three disorders and brain regions (Supplementary Fig. 15G). Ast.C1 showed the greatest number of disease-specific gained open chromatin accessibility in PSP samples in the insula (Fig. 5A, left panel). The significant enrichment of PSP heritability was found in both ast.C1-specific peaks and disease-dynamic peaks (Fig. 5A, right panel and Supplementary Fig. 15F), suggesting ast.C1 is a state harboring PSP heritability-relevant chromatin accessibility, which may contribute to PSP causality. *SOX10* was not only identified as a marker gene for ast.C1, but also exhibited differential activation in PSP (Fig. 5B). We then prioritized functional genes for ast.C1 based on their linked PSP differentially activated CREs associated with GWAS variants (Methods). Surprisingly, we discovered three other SNARE-related genes, *STX7*, *YKT6,* and *STX16,* were activated and involved in SNARE-mediated vesicular transport ^62^ (Supplementary Fig.16A and Supplementary Table 7). The SNARE protein *YKT6* is known to control the stress response and enhance lysosomal activity in Parkinson’s disease ^63^. Notably, *STX7* and other SNARE-related genes, including a well-established PSP risk gene *STX6* ^64^, were differentially upregulated in ast.C1, particularly in PSP (Supplementary Fig. 16B), supporting ast.C1 plays a role in vesicle trafficking. Moreover, other genes involved in the peak-to-gene regulatory circuitry were enriched in endocytosis, regulation of cell migration, synapse and vesicle (Supplementary Table 8). These findings suggest that ast.C1 may enhance intracellular trafficking, including endocytosis and vesicle-mediated transport, in response to tau and myelin accumulation. This process may contribute to maintaining protein homeostasis and preventing cellular dysfunction, potentially mitigating neurodegenerative processes in PSP (Fig. 5C).

In microglia, we observed the highest gain of FTD heritability specifically in mg.C4 (Fig. 5D), implying mg.C4 is a PiD-associated subtype whose accessibility changes might capture FTD causality. Given that mg.C4 involved myelin genes, we tested multiple sclerosis (MS) GWAS as a comparison, but found MS heritability score maximal over a different cluster, mg.C11, marked by genes *TMEM119* and *TREM2* (Supplementary Fig. 15E). Important known FTD disease modifier genes affecting *TDP43* forms of the disease (*GRN*, *SORT1*) were more differentially expressed in mg.C4 than any other microglia type (Supplementary Table 7) ^65^. Interestingly, one CRE in mg.C4 contained a FTD SNP (rs10734151) targeted gene *CTSC*, a capthesin protein which degrades proteins in a lysosome pH dependent manner and was a candidate genome-wide significant hit reported in the original FTD GWAS study, but through a separate locus (rs74977128). We combined CRE with linked FTD GWAS-associated SNPs and performed pathway analysis to define downstream biology. Genes involved in immune system processes, sphingolipid signaling pathway, apoptosis, Fc gamma R-mediated phagocytosis, and regulation of response to endoplasmic reticulum stress were enriched among targeted genes (Tables S9 and S10). Genes differentially activated in PiD in mg.C4 include heat shock proteins and cytokine signaling, suggesting a potential role for regulating microglial pathways related stress and immune response (Fig. 5E). Taken together, mg.C4 might be subject to additional lipid regulation, which enhanced its ER stress response and activated the degradation pathways, including lysosome, phagocytosis, and apoptosis (Fig. 5F).

Additionally, cell-cell interactions (CCIs) inferred by CellPhoneDB based on imputed gene expression (Methods) showed higher CCIs involved in mg.C4 in PiD compared to the other three conditions (Fig. 5G), indicating increased cellular interaction activity in PiD. The cellular interactions orchestrate single-cell functions to maintain homeostasis and regulate physiological processes. We thus honed in on the interactions between mg.C4 and representative neuronal and glial subclusters, such as *RORB*^+^ excitatory neuron C14, the expanded *SST*^+^ inhibitory neuron C9, the largest oligodendrocyte subcluster odc.C7, and PSP-associated ast.C1 (Fig. 5H). We found key ligand-receptor pairs mediating signaling related to myelination, inflammation, and synaptic regulation, including PiD-risk genes *TREM2*, *CLSTN1,* and *PPIA*. *TREM2* plays a protective role in microglia and may promote myelin metabolism ^66,67^. *NOTCH1,* previously identified in the PiD regulatory circuit (Supplementary Table 9), was found to mediate interactions between mg.C4 and neurons via the *CNTN1*∼*NOTCH1* pair, highlighting its role in myelination and neuroinflammation. These results further underscore mg.C4 as a key subcluster responsive to PiD pathology.

## Discussion

Our study presents a detailed analysis of in-house chromatin accessibility and transcriptional profiles of over 600,000 individual nuclei from the matched brain region samples of 41 individuals in tauopathies and controls. We detect cell type and context-specific enhancers by integrating ATAC and RNA nuclei and identify accessibility changes in both pairwise comparison and condition-wide comparison. With the condition dynamic peaks, we interrogate disease GWAS variants, single cell eQTLs, and MPRA functional variants, identify regulatory TF and modules, prioritize target genes, and uncover non-coding circuits disrupted in disease pathology-relevant subtypes.

Our data demonstrates a conserved pattern of chromatin accessibility transition from moderate to high pathology brain regions. There is a general chromatin accessibility closed in astrocytes and inhibitory neurons induced by PiD and PSP, suggesting the mechanisms of over-activation and de-activation responding to disease progression. A common explanation for chromatin accessibility change is that cell type-specific regulatory elements are disrupted by non-coding genetic variants, impacting the gene expression of their linked targets. Integration with disease GWASs shows that accessibility change in disease glia contributes to disease heritability in a cell type-specific way, pinpointing that FTD heritability is highest contributed by dynamic accessibility in microglia. Utilizing the cell type-specific eQTLs and MPRA functional variants, the microglia show significant variant enrichment in dynamic peaks rather than stable peaks, supporting microglia as the FTD target enriched in disrupted accessibility in response to pathology-related and functional variants.

Cell expansion is a common glial feature in neurodegenerative disease. Microglial proliferation and activation may promote cytokine release and contribute to the clearance of pathological proteins in Alzheimer’s disease ^67–70^. We analyzed how subcluster composition changes and identified a significant expansion of two glia subclusters in the insula in individuals with PiD and PSP pathology. These two subclusters, ast.C1 and mg.C4, express genes typically limited to oligodendrocytes, including genes involved in myelin processing genes.

Importantly, we identified ast.C1 as associated with PSP and mg.C4 as associated with PiD, each showing distinct disease-specific roles. Notably, mg.C4 shows the strongest enrichment for FTD heritability and is highly involved in cellular interactions within the PiD context. The CRE module specific to mg.C4 regulated by sn-eQTL and frVars is associated with lysosome function and vesicle-mediated transport. By exploring the non-coding circuit to bridge the gap between genetic variants, regulatory elements, and risk gene expression, we found that in mg.C4, *NOTCH1* consistently exhibits both upregulated gene expression and CRE activation, with its CRE located near PiD-associated genetic variants. *NOTCH1* mediates interactions between neurons and mg.C4 via the *CNTN1*-*NOTCH1* signaling in PiD, where *CNTN1* in neurons is critical for central nervous system myelination ^71^. The binding of *CNTN1* to *NOTCH1* may activate the *Notch* signaling pathway in microglia, modulating their activation states and controlling inflammation ^72,73^. Other target genes in PiD within the non-coding regulatory circuits include lysosome-related gene *CTSC*, as well as genes significantly enriched in pathways including sphingolipid signaling and the regulation of response to endoplasmic reticulum stress (Tables S5, S9 and S10). These findings support the idea that lysosomal pathway activation and immune response in PiD may be driven by lipid-associated stimuli, such as myelin or cellular debris, affecting changes in cell-cell communication to maintain tissue homeostasis.

In parallel, we revealed a PSP-associated myelin-related astrocyte, ast.C1, in the insula, where epigenomic integrity was preserved with cell expansion, suggesting a protective role. Chromatin accessibility in this expanded astrocyte subtype is uniquely open in PSP compared to controls, and both subcluster-specific and PSP-upregulated peaks capture PSP heritability. PSP risk variants are enriched in PSP up-regulated dynamic enhancers of SNARE proteins, particularly *STX7*, whose expression is also increased. Top regulatory transcription factors show altered accessibility in PSP glia, with *NFIX* significantly targeting a tau-binding gene, *HSPA2*, potentially influencing *MAPT* upregulation (Supplementary Fig. 15B). Moreover, *SOX10*, differentially activated in PSP, specifically targets another SNARE protein, *STX4*. Integrating genetics, epigenomics, and expression data, we found PSP-relevant non-coding circuits involve SNARE interaction, synapse, and vesicle (Supplementary Table 8). Building on this, we propose that the expanded astrocyte subtype, ast.C1, may facilitate the degradation of misfolded proteins by promoting increased autosome-phagosome fusion via SNARE proteins.

In summary, we provide a cross-disorder atlas linking gene regulation, chromatin dynamics and cellular functions across three tau disorders, to highlight disorder-specific glial states of differential resilience. We reveal epigenomic dynamics and map genetic variants to their target through CREs, enhancing our understanding of disease regulatory circuits. We provide molecular targets linked to polygenic disease risk, prioritizing genes for experimental validation to inform causal mechanisms and therapeutic strategies. Our data enhance the understanding of glial contributions to various tauopathies at the single-cell level and underscore the importance of cross-disorder, cell-specific chromatin profiling in brain regions with moderate pathology.

## Methods

### snATAC-seq data processing

The nuclei extraction and sequencing setting of snATAC-seq data matched those described in our previous published work ^19^. Using ArchR version 1.0.3 ^29^, we first retained cells with TSS enrichment >= 2 and a minimum of 1,000 fragments per sample. Additionally, we re-filtered out low-quality cells for 14 samples with sample-specific stringent cutoffs based on QC plots (Supplementary Fig. 1). Doublets were removed using the filterDoublets function. Dimensionality reduction was performed using iterative Latent Semantic Indexing with six iterations and 25,000 variable features. We applied the Harmony batch effect correcting for preparation batches (Supplementary Fig. 2C). Clustering was conducted using the addClusters function with a resolution of 0.1, generating 10 main clusters. Marker gene scores for each cluster were calculated using the getMarkerFeatures function, retaining those with an FDR <=0.01 and log2 fold change (Log2FC) >= 1.25. Canonical marker genes were used to annotate the main brain cell type for each cluster, and two small undefined clusters were filtered out. Finally, we integrated the data with our in-house snRNA-seq dataset ^19^ using the addGeneIntegrationMatrix function, ensuring the consistency of cell type annotation (Supplementary Figs. 2D-E). For the downstream analysis, we removed one PreCG sample and one insula sample of the PiD case P2301, defined as an outlier in our previous study ^19^. Based on differences in tissue microdissection during sample preparation where white matter was grossly removed from PreCG and not insula samples, the cluster composition of samples exhibited regional heterogeneity and showed greater diversity between brain regions than between diseases (Supplementary Fig. 13A). Neurons were more frequently found in PreCG, with an average cell percentage of 42.6% ranging from 35.2% to 62.2%, and the highest occupation in 67.5% of samples (27 of 40). In contrast, cell proportions of control samples were more stable than disease samples, with 7 out of 9 control samples displaying the highest proportions in neurons. In the mid-insula, oligodendrocytes emerged as the predominant cell type in nearly all cases (94.7%, 36 of 38), with an average of 49.0%, ranging from 34.3% to 66.1%.

## Subcluster identification

We performed iterative LSI and clustering with different resolutions using Louvain to construct subclusters for each cell type. The resolution, which generates clusters with meaningful cell numbers, clear boundaries, and obvious identified markers, was selected. We examined the cell proportions among library batches, sequencing batches, and samples (Supplementary Fig. 8). We removed subclusters with less than 100 cells, as well as those contributed by one batch or one sample uniquely (ast.C6, mg.C8, mg.C10, mg.C15). We end up identifying 57 subclusters in total. To confirm the corresponding cell type of each subcluster, we calculated marker gene scores across all subclusters using ArchR’s getMarkerFeatures function, filtering by FDR < 0.1, and examined the Log2FC of canonical cell type markers (Supplementary Fig. 9B and Supplementary Table 11). In addition to the raw cell set, we created a high-quality cell set consisting of cells with a TSS enrichment score ≥ 4 to replicate the identification (Supplementary Table 12). Subclusters that did not express the corresponding cell-type markers in both datasets (Log2FC > 1) were filtered out. We then removed ambiguous subclusters that highly express markers of more than one cell type. Some potential hybrid clusters were retained when another cell type marker was expressed, but not as the highest. The average gene scores of each subcluster show consistent cell type marker expression (Supplementary Fig.9B). We identified 50 subclusters, including 10 astrocytes, 10 microglia, 12 neurons, 8 oligodendrocytes, and 10 OPC subclusters.

## Subcluster Annotation

We used marker gene scores identified across subclusters to confirm the cell type identity of subclusters, and marker gene scores identified within each cell type to assign subtype functional categories. To understand the functional diversity among subclusters. We first assigned functional groups based on the significant upregulation of disease signature genes. Next, we determined the marker gene scores within the same cell type (Supplementary Table 13) and conducted enrichment analysis for markers using enrichR. The enriched GO terms (FDR < 0.1) and the associated genes were selected as the functional description for each subcluster relative to the other subclusters within the same cell type.

For astrocytes, *GFAP* was widely expressed in all subclusters, with the lowest in ast.C1 and the highest in ast.C5, based on the marker gene scores identified across all subclusters (Supplementary Fig. 9B). The hierarchical clustering of markers called within astrocytes reveals the four main categories of astrocytes (Figs. 4B-C). Reactive astrocytes, represented by ast.C5, ast.C7, ast.C8, and ast.C9, highly expressed *GFAP*, with slight variations in the marker peaks (Fig. 4K). Ast.C2 was related to synaptic transmission. Ast.C3 was related to antigen presentation, exhibiting marker peaks of *HLA-DMB* (Figs. 4C and 4K). Both ast.C1 and ast.C4 were myelination-related and oligodendrocyte like, with ast.C1 displaying prominent marker peaks for *KLK6* and *PLP1* (Fig. 4 K). Ast.C10 was linked to inflammation and had the lowest MAPT gene score, and ast.C11 was metabolism-related (Fig. 4C). When compared with other astrocyte subclusters, ast.C1 highly expressed oligodendrocyte markers, including *MOBP*, *TF*, and *OPALIN*, as well as disease signature genes found in oligodendrocyte, such as *CNTN2*, *SLC5A11* (Fig. 4 K and Supplementary Fig. 9B).

For microglia, we identified a total of 10 subclusters, including two homeostatic states mg.C16 *NAV*^+^, mg.C14 *P2RY13*^+^*CTSS*^+^; two transitional state microglia with both homeostatic and inflammatory marker activation: mg.C11 *TMEM119*^+^*TREM2*^+^, mg.C13 *C1QA*^+^*P2RY12*^+^*TREM2*^+^; one disease-associated microglia (DAM) mg.C12 *APOE*^+^*TYROBP^+^*; two myelination-related microglia mg.C4 and mg.C6 (*SOX10*^+^*PLP1*^+^); two synaptic transmission genes enriched states mg.C3 and mg.C7 (*SNCB*^+^*DRD4*^+^), and a T cell activation gene enriched subcluster mg.C9 *CD247*^+^*OPTN*^+^. The later five subclusters, except C6, also express autophagy markers *BPTOR* and *OPTN* (Figs. 4E and 4G). Marker peaks were consistently up-regulated in specific subclusters, such as *PLP1* and *SORT1* in mg.C4, *RIN3* in mg.C13, *CD48* in mg.C9, *TREM2* in mg.C13 and mg.C11, and *LGALS3* in mg.C7 and mg.C14. Compared with mg.C4, mg.C6 exhibited greater chromatin accessibility around *RIN3*, *CD48,* and *TREM2* (Fig. 4J).

All of the 8 oligodendrocyte subclusters are up-regulated genes related to myelination, such as *CNTN2*, *OPALIN*, *CARNS1*, *PLP1*, and *OLIG1* (Supplementary Fig. 9B). Furthermore, 4 of these subclusters (odc.C3, odc.C4, odc.C5, odc.C7) also up-regulated *SLC5A11*, which was associated with lipid transport. Odc.C7 was also inflammatory oligodendrocytes enriched in cytokine receptor binding, and odc.C8 was stress-related with markers enriched in DNA repair. In addition, we annotated 10 subclusters for OPC, three of which up-regulated AD signature genes. Opc.C1 expressed *IL6R*, which was associated with immune response, whereas opc.C8, opc.C9, and opc.C10 expressed *GALR1*, associated with neuronal activity. OPC.C9 and OPC.C10 also expressed *LRP1* related to lipid transport (Supplementary Fig. 9B). For neurons, we identified four inhibitory neurons and eight excitatory neurons (Supplementary Fig. 9B). The four inhibitory neurons consist of four *SLC32A1*^+^ *GAD1*^+^ cells. Among them, three expressed additional markers: *PVALB*^+^(neu.C7), *SST*^+^ (neu.C8) and *ADARB2*+ *VIP*+ (neu.C6). The four excitatory neuron subclusters expressed layer-specific markers: neu.C5 and neu.C11 were *FEZF2*^+^, neu.C14 was *RORB*^+^, and neu.C13 was *CUX2*^+^.

## Marker comparison of microglia subclusters with a public dataset

We obtained the complete list of marker genes of microglial subtypes from ^55^ and compared them with the marker gene scores of microglial subclusters defined in our study. We computed the pairwise Jaccard scores and assessed the significance of the overlap using Fisher’s exact test. P values were adjusted for multiple testing using the Benjamini–Hochberg correction. The results, including Jaccard scores and significance levels, were visualized in a heatmap.

## CRE identification and validation

We identified potential cis-regulatory elements in each subcluster under each condition based on peak-to-gene relationships and peak-to-peak co-accessibility using ArchR. We used the addCoAccessibility function to calculate co-accessibility and the addPeak2GeneLinks function to calculate peak-to-gene links. A gene’s promoter peak was defined when the Pearson correlation coefficient between peak accessibility and gene expression was greater than 0.45 (FDR < 0.1). An enhancer peak was defined as any peak that was not a promoter peak but whose accessibility correlated with that of a promoter peak (Pearson coefficient > 0.5 and FDR < 0.1). To validate our candidate enhancers, we collected reference enhancers from 11 public resources, including ENCODE ^30^, activity-by-contact (ABC) model predictions ^31^, FAMTOM5 (Functional Annotation of Mammalian Genomes 5) ^32,33^, and three single-cell studies ^17,34,35^. We downloaded the peak sets from the ENCODE portal (https://www.encodeproject.org) with the following identifiers: ENCFF198KYT, ENCFF791URB, ENCFF395QLP, and ENCFF815WRM. We classified a CRE as "known" if it overlapped with the loci of any reference enhancers and as "novel" if it did not.

## Identification of differentially accessible regions

Using ArchR’s iterative peak calling strategy, we generated pseudobulk replicates and identified peaks for each subcluster, then created a consensus peak set. To identify differentially accessible regions between disease and control within each cell type and region, we performed differential accessibility testing using the Wilcoxon test in the discovery dataset. To ensure the robustness of DAR calling, we performed downsampling 10 times with 30 nuclei per sample for each cell type and re-conducted the differential test using the Wilcoxon test. The significance cutoff for the discovery and downsampled datasets was set at P-value < 0.001 and |log2 fold change|>= 1.2.

## Identification of dynamic and stable peaks per cell type

We first identified cell type-specific peaks within the consensus peak set. We labeled each peak as 1 (present) or 0 (absent) to indicate its presence in each subcluster based on the original peak calling results. These labels were then concatenated to determine peak presence across each cell type. Next, we identified marker peaks for each of the four conditions within each subcluster by comparing one condition against the other three conditions. Cell type-specific disease marker peaks were determined based on P-value < 0.05 and |log2 fold change| > 1.

## Heritability partition by LDSC

Integrating with GWAS studies of AD, PiD, PSP, and MS ^64,76–78^, we applied stratified linkage disequilibrium score regression (S-LDSC) to partition trait heritability. We used two metrics to measure the contribution of a functional category C to trait heritability: enrichment and standardized effect size r* ^79–81^. The enrichment score evaluates whether the per-SNP heritability r_C_ is greater in the category than overall r:

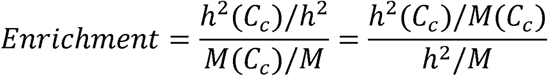

where h^2^is the heritability, and M is the number of SNPs; C represents the category.

Since r_C_ depends on trait heritability and the size of the annotation, the standardized effect size (τ*) is used to compare r_C_ between different traits or annotations.

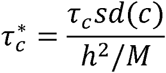

We applied LDSC to multiple peak sets, including subcluster-specific peaks, cell type-specific dynamic and stable peaks, dynamic peaks classified as upregulated or downregulated within each disease, and peaks overlapping with transposable elements.

## Dynamic CREs Associated with GWAS Variants

To identify regulatory circuitry mediated by GWAS genetic variants, we linked CREs with GWAS SNPs based on the following criteria: (1) SNPs localized within CREs ± 2kb upstream or downstream, and (2) CREs containing a non-GWAS SNP in linkage disequilibrium (LD) with a GWAS SNP. LD scores were calculated by plink ^82,83^ with variants in Europeans (1000 Genomes Phase 3). CREs that exhibited dynamic changes (|log2FC| > 1) in disease conditions were selected. Genes with dynamic CREs associated with GWAS SNPs (either directly or through LD, R² > 0.8) were used for functional enrichment analysis by ShinyGO 0.80 ^84^.

## Reproducing dynamic peaks analysis using SEA-AD snATAC-seq

We used publicly available SEA-AD snATAC-seq data from the middle temporal gyrus to replicate findings of dynamic peaks. The SEA-AD dataset includes 24 pre-defined cell subclasses across four conditions: no AD, low AD pathology, intermediate AD pathology, and high AD pathology. We consolidated these subclasses into six major cell types: astrocytes, microglia, oligodendrocytes, OPC, excitatory neurons, and inhibitory neurons, while excluding VLMC and endothelial cells. Utilizing the 218882 consensus peaks defined in SEA-AD, we first identified cell-type marker peaks by calling marker peaks within each subclass and then aggregating them to the cell-type level. Marker peak calling was performed using Scanpy with FDR < 0.1 and log2 fold change > 1. Subsequently, we identified marker peaks for each of the four conditions within each subclass (FDR < 0.1). We selected condition marker peaks that were also identified as cell-type marker peaks to define cell-type dynamic peaks. Peaks that remained unchanged across all four conditions were classified as stable peaks. Finally, we performed LDSC enrichment analysis using AD GWAS data ^85^ to partition AD heritability among cell-type dynamic peaks, stable peaks, and dynamic peaks identified in each condition.

## Massively parallel reporter assay

We performed a lentiviral-based Massively Parallel Reporter Assay (lentiMPRA) to functionally assess the transcriptional regulatory activity of candidate sequences. Oligonucleotides containing each test sequence upstream of a minimal promoter and a barcoded reporter were cloned into a lentiviral backbone. The resulting plasmid library was packaged into lentivirus and transduced into the human microglial cell line HMC3 (ATCC CRL-3304), with an estimated coverage of 100 barcodes per candidate sequence and approximately 100 integrations per barcode. After overnight incubation, the virus was removed, and cells were cultured for an additional two days to allow sufficient expression of the reporter gene. Genomic DNA and total RNA were then extracted, and barcoded reporter transcripts were quantified by high-throughput sequencing. Parallel DNA sequencing of the integrated barcodes was used to normalize for integration efficiency. Regulatory activity was calculated as the RNA/DNA ratio for each barcode, averaged across barcodes per sequence. Functional variants were defined as those showing significantly different regulatory activity between reference and alternative alleles, with a false discovery rate (FDR) below 0.05.

## MPRA and single-nucleus eQTL enrichment analysis

We analyzed the distribution of P values, FDR, and log2 fold changes across all MPRA variants, identifying functional MPRA variants with a P value < 0.05. We obtained the single nuclei eQTLs from ^52^ with an FDR < 0.00, and from ^86^ with significant_by_2step_FDR column label as Yes. To assess the enrichment of MPRA frVars and significant eQTLs, we used Fisher’s Exact Test to examine variants overlapping dynamic and stable peaks. An eQTL was considered colocalized with a peak if the eGene matched the gene linked to the peak. We performed variant enrichment analyses in dynamic and stable peaks across different peak types: all peaks, CRE and gene body peaks, CRE peaks alone, and enhancer-only peaks. For genes with differentially accessible enhancers containing MPRA frVars across disease conditions, we conducted functional enrichment analysis using ShinyGO 0.80 ^84^, selecting enriched terms with an FDR-adjusted P value < 0.1.

## Detection of the CRE module linked with frVars or eQTLs

To define co-occurring regulatory modules based on chromatin accessibility across subclusters, we first generated 823 pseudobulked samples, considering two to five replicates per disease within each of the 50 subclusters, using the ArchR strategy. For each peak in each pseudobulk sample, we aggregated peak counts, depth-normalized them, and applied a log2 transformation. The resulting peak-by-pseudobulk matrix was then quantile normalized across samples. Next, we calculated the average peak activity per peak at the subcluster level, and subsetted the matrix for CRE peaks overlapping functional variants. Hierarchical clustering identified co-occurring CRE modules with their subcluster specificity visualized in heatmap. TF regulators of modules were identified by overlapping predicted TFBS of CREs with TF enrichment analysis using MEME.

## Identification of regulatory TFs and their target genes

To determine if a TF motif plays a potential regulatory role, we first calculated the per-cell accessibility for each TF motif using ChromVar ^87^, and selected candidate motifs with quantile z-scores greater than 0.5. Subsequently, we calculated the Pearson correlation between TF activity and TF expression across subclusters, identifying motifs with correlation coefficients greater than 0.5 (FDR < 0.1). This process led to the identification of 164 TF motifs with regulatory roles. Additionally, we identified the top regulatory motifs whose ChromVar variability z-score is above the 90th quantile. To explore the TF variability changes in disease, we calculated delta variability between disease and control (delta z-score) and identified the top 10 TFs with the highest delta z-score for each cell type.

We calculated case-control differentially activated TFs for each cell type and subcluster using the ArchR’s getMarkerFeatures function with the Wilcoxon test on the gene score matrix, retaining significant genes with FDR < 0.1. To identify putative target genes for a TF, we first selected genes whose gene scores are positively correlated with TF activity (Pearson coefficient >0.25 and FDR <0.1). For each gene, we calculated gene-TF linkage scores based on the peaks that are linked to the gene and contain the motif ^88^: Within each cell set (e.g., a disease subcluster like PSP ast.C1), we selected peaks containing the motif that also had accessibility positively correlated with gene activity (Pearson coefficient > 0.1 and FDR <0.1). The linkage score was calculated as the aggregated square of Pearson coefficients of all selected peaks. Finally, we identified genes with linkage scores above the 80th quantile as potential target genes within the given cell set.

## Prediction of TF binding sites in subcluster regulatory elements (in ast.C1 and mg.C4)

To identify TF targets by mediating binding events in accessible chromatin, we predicted TF binding sites on the marker peaks of subclusters using TOBIAS ^89^, as described in our previous study ^19^. We used the TOBIAS functions *ATACorrect* to correct Tn5 insertion bias in mapped ATAC-seq reads and then utilized *FootprintScores* to calculate foot printing scores across peak regions. To estimate the binding positions of individual TFs on marker peaks, we applied *BINDetect* which combines the predicted footprint scores with TF binding motif information. Genes with CREs bounded by any TF were considered as targets of that TF for further analysis.

## Cellular composition change analysis

To infer composition changes between disease and control groups, we employed a linear regression model by Limma ^90^ to assess the statistical significance of compositional changes between disease and control groups while adjusting for covariables such as age and PMI. A p-value threshold of <0.05 was applied to determine significance. To replicate the cellular changes in ast.C1 and mg.C4, we performed a bootstrapped subcluster composition analysis. Over 15 iterations, we sampled 20% of cells from the entire dataset and computed subcluster proportions for each condition. We then applied the Wilcoxon rank-sum test (Wilcox. test in R v4.2.2) to compare composition differences between disease and control groups. Bootstrapped p-values were adjusted for multiple testing using the Benjamini–Hochberg method.

## Analysis of single-nucleus RNA-seq-derived subclusters

To validate the snATAC-seq-derived subclusters ast.C1 and mg.C4, we aligned them with pre-identified subclusters from in-house snRNA-seq data. We first aligned the snATAC-seq dataset with the snRNA-seq dataset to determine the mapped RNA subgroup and then identified markers for the mapped subgroup to confirm their consistency with the ATAC subcluster. We restricted the cellular alignment within each cell type and subset both the ATAC and RNA datasets to cell type level using Seurat’s farmwork ^91^. The snATAC-seq gene activity and snRNA-seq gene expression were normalized separately. Canonical correlation analysis (CCA) was then performed to identify anchors for pairs of cells using Seurat’s FindTransferAnchors function. We retained results with a prediction score greater than 0.5. We found that 100% of mg.C4 and 89 % of mg.C6 mapped to an RNA subgroup insula-microglia-3, and 86% of ast.C1 mapped to insula- astrocyte-3 (Supplementary Fig. 11B). After alignment, the cellular composition changes in mapped RNA subclusters were measured using bootstrapping with 15 iterations with Wilcoxon rank-sum test.

## Epigenomic stability analysis with single-nucleus methylation data

To assess disease-related epigenomic stability at subcluster level, we measured the extent to which disease chromatin accessibility changes occurred at cell type-specific hypermethylated regions in each subcluster, as described in our previous study ^19^. First, we categorized cells into four conditions (Control, AD, PiD and PSP) within each subcluster, and called peaks for each condition using MACS2 in ArchR. Second, we integrated public single-nucleus methyl-3C sequencing data ^92^ to quantify methylation levels for each ATAC peak. We used ALLCools ^93^ functions allc-to-region-count to calculate methylation level and generate-dataset to compute the hyper-methylated score. ATAC peaks were considered methylated if they met the criteria of a hyper-methylation score >= 0.9 and a methylation level > 0. Third, we performed a permutation test (n= 10,000) to compare the proportion of chromatin-accessible regions annotated as methylated between disease and control groups. A higher proportion of methylated regions in disease samples suggests a loss of heterochromatin, while a higher proportion in control samples indicates heterochromatin gain in disease, both reflecting epigenomic relaxation or instability. Empirical p values were adjusted for multiple testing (*** FDR ≤ 0.001, ** FDR ≤ 0.01, * 0.01 < FDR ≤ 0.05).

## Cell-cell interactions analysis

To infer cellular interactions across conditions, we used the CellphoneDB DEG-based method. First, we imputed expression values for cells identified from snATAC-seq data for each individual. The snATAC-seq data were processed using the Signac package ^94^, and alignment with snRNA-seq data was performed using Seurat’s FindTransferAnchors function, employing canonical correlation analysis (CCA). We downsampled each sample to 35% of its total cell count while maintaining the subcluster distribution. For each sample, we calculated the targeted number of cells representing 35% of the total, then randomly sampled from each subcluster in proportion to its original representation within the sample. This approach preserved the subcluster structure after downsampling. Differentially expressed genes (DEGs) were calculated on the downsampled dataset using imputed expression values with Seurat’s FindMarkers function. DEGs for each disease condition were identified through pairwise comparisons with the control samples, while control samples were collectively compared against all three disease conditions. DEG analysis was conducted separately for each cell type and subcluster, retaining up-regulated genes with a log2 fold change (log2FC) > 0.25 and a false discovery rate (FDR) < 0.1. Finally, we ran CellphoneDB for each of the four conditions using default parameters to infer cell-cell interactions based on the identified DEGs. The differences in the number of cellular interactions between conditions were assessed using a chi-square test, with a false discovery rate (FDR) threshold set at < 0.05.

## TF enrichment

TF motif enrichment was carried out using the runAme function from MEME Suite in R. For motif enrichment in cell condition-dynamic peaks compared to stable peaks within each cell type, we used the JASPAR 2022 non-redundant motif database ^95^. Other peak sets were analyzed using the CISBP 2.00 Homo sapiens database ^96^. Significantly enriched functional terms and TFs were retained with an FDR < 0.1.

## RNAscope and immunofluorescence

Experimental validation was performed using RNAscope and immunohistochemistry (IHC) on formalin-fixed paraffin-embedded (FFPE) human insular brain tissue from one PSP case (sample ID: I1_5) and one control case (sample ID: i3_5_at). Brain tissue was sectioned at a thickness of 6µm. RNAscope™ assays were performed using an automated Leica Biosystems platform at the UCLA Translational Pathology Core Laboratory. Target RNA-specific oligonucleotide probes used for mRNA of human *SOX10* (CATALOG # 484128-C4) and *PLP1* (CATALOG# 499278) were synthesized by Advanced Cell Diagnostics. GFAP protein was detected by IHC using anti-GFAP antibody (Dako, clone M0761) at a 1:200 dilution. IHC staining was performed after completing the RNAscope procedure. Images were analyzed using Phenochart 2.2.0 with spectral unmixing to minimize background signals. Signals located within or adjacent to nuclei and exceeding the average background intensity were considered positive RNAscope signals.

## Acknowledgements

We thank Lawrence Chen, Mai Yamakawa, and Sen Ma for technical support and coding assistance; Connor Webb for human tissue requesting; and all lab members for valuable comments. This work was supported by NIH grants (R01 AG075802 [J.E.R. and L.T.G.], R01AG068030 [K.J.B], RF1AG065926 [K.J.B], R01AG050986 [K.J.B], R01ES033630 [K.J.B]).

The UCSF Neurodegenerative Disease Brain Bank is supported by NIH grants AG023501 and AG019724, the Rainwater Charitable Foundation, and the Bluefield Project to Cure bvFTD.

## Author Contributions

X.H. and J.E.R. designed the study. X.H. performed all computational analysis, interpreted results, and wrote the manuscript. T.Z. analyzed RNAscope and IHC data. T.R. and J.L. performed MPRA experiments under the guidance of T.R. and K.J.B. W.W.S., L.T.G., A.N.L., and S.S. performed regional neuro-pathological assessments and scoring. J.E.R supervised the entire study, reviewed and edited the manuscript.

## Ethics declarations

The authors declare no competing interests.

## Supplementary information

Supplementary Fig. 1. Quality control plot for 14 samples with sample-specific thresholds.

Supplementary Fig. 2. Cellular abundance and gene signatures among cell types.

(A-B) UMAP plots displaying gene scores (A) and imputed expression (B) for canonical markers of each cell type.

(C) UMAPs of cells colored by sequencing and library batches.

(D) Alignment between snATAC-seq and in-house snRNA-seq showing the percentage of ATAC cells (rows) mapped to RNA cell types (columns), performed using Seurat’s Canonical Correlation Analysis (CCA).

(E) Bar plot showing the number of cells detected per cell type in the snATAC-seq and snRNA- seq datasets.

Supplementary Fig. 3. TFs with regulatory roles across cell types and diseases.

(A) Prioritization of TFs with regulatory roles across subclusters. The x-axis shows the Pearson correlation between TF activity and TF expression, and the y-axis shows TF variability defined by ChromVar. Top TFs with a z-score deviation above the 90th percentile and a correlation coefficient > 0.5 between activity and expression are highlighted in red (left). UMAP plots on the right depict the gene score, variability z-score, and gene expression of selected TFs.

(B) The delta TF variability for disease and control groups.

Supplementary Fig. 4. Characterization of CREs across genomic features, reference resources and diseases.

(A) Pie chart depicting the distribution of consensus peaks across genomic contexts (CRE, promoter, intronic, exonic, or distal regions) before and after CRE identification.

(B) Bar plot showing the percentage of snATAC-seq-derived enhancers that overlap with reference enhancers from various resources.

(C) Bar plot showing the number of identified consensus peaks distributed across cell types.

(D) Bar plot displaying the number of dynamic and stable peaks detected in each subcluster.

(E) Bar plots depicting the distribution of consensus peaks and DARs across genomic contexts (CRE, promoter, intronic, exonic, or distal regions).

(F) Pie chart of differentially accessible enhancers categorized as known or novel.

(G) Bar plots showing the number of differentially accessible CREs in glia and neurons (top) and glia-specific CREs by glia types (bottom).

(H) Bar plot displaying the number of differentially accessible CREs detected in the downsampled dataset, categorized into up-regulated and down-regulated peaks.

Supplementary Fig. 5. Additional analysis of dynamic peaks across diseases.

(A) Pie charts showing the distribution of GWAS SNPs located in dynamic peaks across conditions. (B) AD heritability enrichment in dynamic and stable peaks within microglia in the SEA-AD dataset.

Supplementary Fig. 6. Additional analysis of MPRA.

(A) Distribution of p-value, FDR, and effect size for all MPRA variants (left), MPRA variants in dynamic peaks (middle), and stable peaks (right).

(B) Bar plot displaying the number of cell type-specific peaks containing MPRA frVars, with peaks categorized by gene context (distal, enhancer, promoter, exonic, or intronic).

(C) Heatmap showing regulatory modules of CREs with eQTLs across microglial subtypes. Peak accessibility in pseudobulked samples was log2-transformed after depth normalization, and the mean values of subcluster were quantile normalized. Functional enrichment of CRE-linked genes was analyzed using enrichR.

(D) TF-target network of module 10 driven by nine key TFs was specifically activated in mg.C4, targeting CREs linked to genes enriched in vesicle-mediated transport and lysosomal functions. CREs linked to target genes colored according to their maximal differential accessibility score, calculated as -log(P-value) x |log2FC| from marker peak calling.

**Supplementary** Fig. 7**. Key TF regulators of module10 drive CRE activation in mg.C4.** Heatmaps show the normalized chromatin accessibility of CREs bound by the corresponding TF across microglial subclusters, with CRE-linked genes displayed. Peak accessibility in pseudobulked samples is log -transformed after depth normalization, and the mean values for each subcluster are quantile normalized.

**Supplementary** Fig. 8**. Identified subclusters with negligible batch effects.** UMAPs are colored by subclusters, sequencing batches, and library batches for each cell type. The proportions of cells per subcluster contributed by different batches, brain regions, conditions, and genders are displayed on the right.

Supplementary Fig. 9. Cell percentages and gene signatures of identified subclusters in human brain.

(A) Bar plot displaying the number and percentage for subclusters, labels representing cell number, subcluster frequency within the cell type, and overall subcluster frequency.

(B) Gene signatures of 50 subclusters, defined by marker gene scores compared across all subclusters.

(C) UMAPs showing subclusters and *SOX10* gene scores in astrocytes and microglia.

Supplementary Fig. 10. RNAscope ISH for *SOX10* and *PLP1* combined with GFAP IHC in human insular tissue from a PSP patient and a control. Representative images show depression and co-localization of *SOX10* and *PLP1* mRNA, with both signals detected in a subset of GFAP-positive astrocytes. (Scale bar: 50 µm)

Supplementary Fig. 11. Reproducing of cell proportion changes in ast.C1 and mg.C4.

(A) MAPT gene activity and imputed expression in microglia (top) and astrocytes (bottom) derived from snATAC-seq.

(B) Percentages of snATAC-seq astrocytes (top) and microglia (bottom) mapped to the pre- defined snRNA-seq subtypes. Cellular alignment was performed using Seurat CCA with prediction scores > 0.5.

(C) Boxplots showing cell composition in each condition for the snATAC-seq subclusters (top) and the mapped snRNA-seq subtypes (bottom). Composition differences between disease and control groups were assessed using the Wilcoxon test after bootstrapping 15 times. FDR thresholds: *< 0.05; ** <0.005; *** < 0.001.

Supplementary Fig. 12. Microglia Subtypes alignment with sn-RNAseq studies.

(A) Overlap of marker genes with microglia subtypes from Schirmer et al ^54^ by Jaccard score and Fisher’s exact test. Significant overlaps with FDR-adjusted p value < 0.1 are marked with asterisks.

(B) UMAP plots of microglia subclusters profiled in snRNA-seq (top left) and snATAC-seq (top right).

(C) UMAPs display selected marker genes expressed in snRNA-seq-derived microglia subclusters.

Supplementary Fig. 13. Cell distribution across subclusters in samples and changes in cell type composition.

(A) Distribution of subclusters across samples, split by disease, with sample IDs colored by brain regions

(B) Boxplots displaying the relative abundance of each cell type across conditions. Changes in cell composition between disease and control were modeled using linear regression via Limma, adjusting for age and PMI. *P < 0.05.

**Supplementary** Fig. 14**. Subcluster composition changes in disease vs. control.** Boxplots displaying the relative abundance of each subcluster across conditions. Changes in cell composition between disease and control were modeled using linear regression via Limma, adjusting for age and PMI. *P < 0.05.

**Supplementary** Fig. 15**. Disease heritability partition by LDSC in subcluster-specific peaks.** (A-B) Target genes *SOX10* (A) and *NFIX* (B) under different conditions in ast.C1. The x-axis represents the Pearson correlation between motif activity and gene activity, and the y-axis indicates gene-TF linkage scores.

(C) Bar plot showing astrocyte-specific heterochromatin remodeling in PSP. The values represent the percentage of ATAC peaks annotated as hypermethylated (mC%) and the absolute difference between PSP and control. A higher mC% in PSP indicates a loss of heterochromatin, while a lower mC% indicates a gain of heterochromatin. Methylation was quantified using single- nucleus methyl-3C sequencing data ^76^ with ALL Cools ^77^. Empirical P-values were obtained from a permutation test (n = 10,000) and corrected for multiple testing (*** FDR ≤ 0.001, ** FDR ≤ 0.01, * 0.01 < FDR ≤ 0.05)

(D) FTD GWAS heritability partition in OPC subcluster-specific peaks measured by LDSC enrichment.

(E-F) GWAS heritability partition of MS in microglia (E) and PSP (F) in neuron subcluster- specific peaks, using both LDSC standardized effect size and enrichment metrics. (significant for r*, FDR *< 0.05; ** <0.005; *** < 0.001).

(G) Number of DA-CREs detected at the subcluster level. Bar plots show the distribution of DA- CREs for each disease in each subcluster, with upregulated DA-CREs in red and downregulated DA-CREs in green. The significant cutoff was set at |log2fc| >= 1.2 and FDR< 0.1. Ast, astrocytes; neu, neurons; MG, microglia; ODC, oligodendrocytes.

**Supplementary** Fig. 17**. SNARE interaction involved in PSP ast.C1.** (A) Chromatin accessibility track illustrating a PSP-upregulated *STX7* enhancer near PSP-associated variants. (B) Differentially expressed genes (FDR-adjusted p value <0.1) in ast.C1 involved in SNARE interactions.

Supplementary Table 1. Differential cellular composition by control vs. diagnosis group in each brain region.

Supplementary Table 2. Transcription factor enrichment in subcluster marker peaks. Supplementary Table 3. *SOX10* target genes shared or unique among ast.C1, mg.C4, and odc.C7. Supplementary Table 4. Predicted target genes of transcription factors based on TF binding sites. Supplementary Table 5: Functional enrichment of *SOX10* target genes in subclusters.

Supplementary Table 6. Significant differential *MAPT* activity between disease and control in subclusters (FDR < 0.1).

Supplementary Table 7. Dynamic peaks in ast.C1 (|log2fc|>1) associated with PSP GWAS SNPs. Supplementary Table 8. Functional enrichment of genes with CREs altered in PSP in ast.C1 and associated with PSP GWAS.

Supplementary Table 9. Dynamic peaks in mg.C4 (|log2fc|>1) associated with FTD GWAS SNPs. Supplementary Table 10. Functional enrichment of genes with CREs altered in PiD in mg.C4 and associated with FTD GWAS.

Supplementary Table 11. Marker gene scores across all unfiltered 57 original subclusters in the raw cell set.

Supplementary Table 12. Marker gene scores across all unfiltered 57 original subclusters in the high-quality cell set (TSS ≥ 4).

Supplementary Table 13. Marker gene scores for subclusters compared to all other subclusters within each cell type.

