## Supplemental Figs for "Single-nucleus epigenomic dysregulation unmasks genetic risk-associated neurodegenerative glia states"

### Supplementary Fig. 1

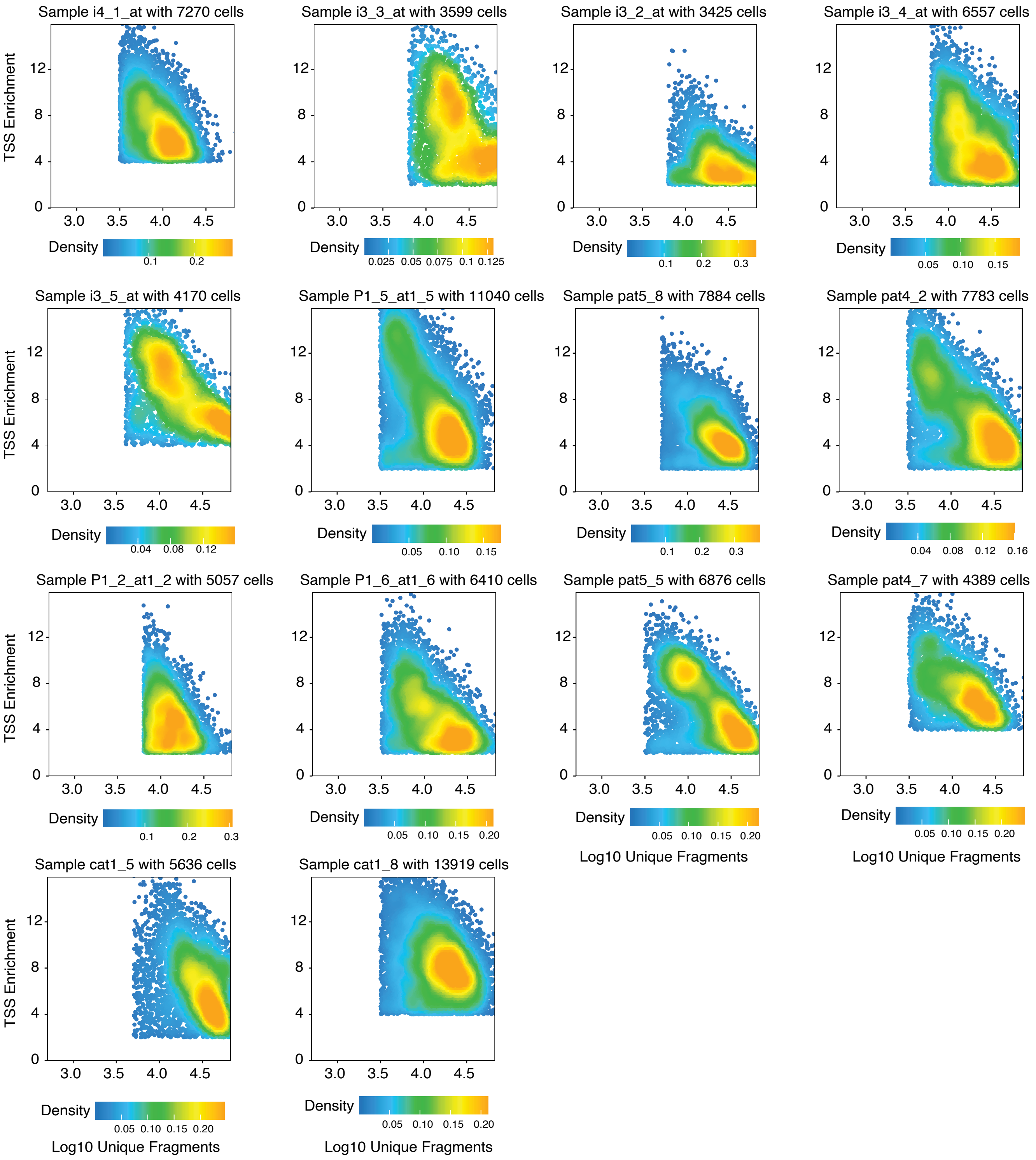

### Supplementary Fig. 2

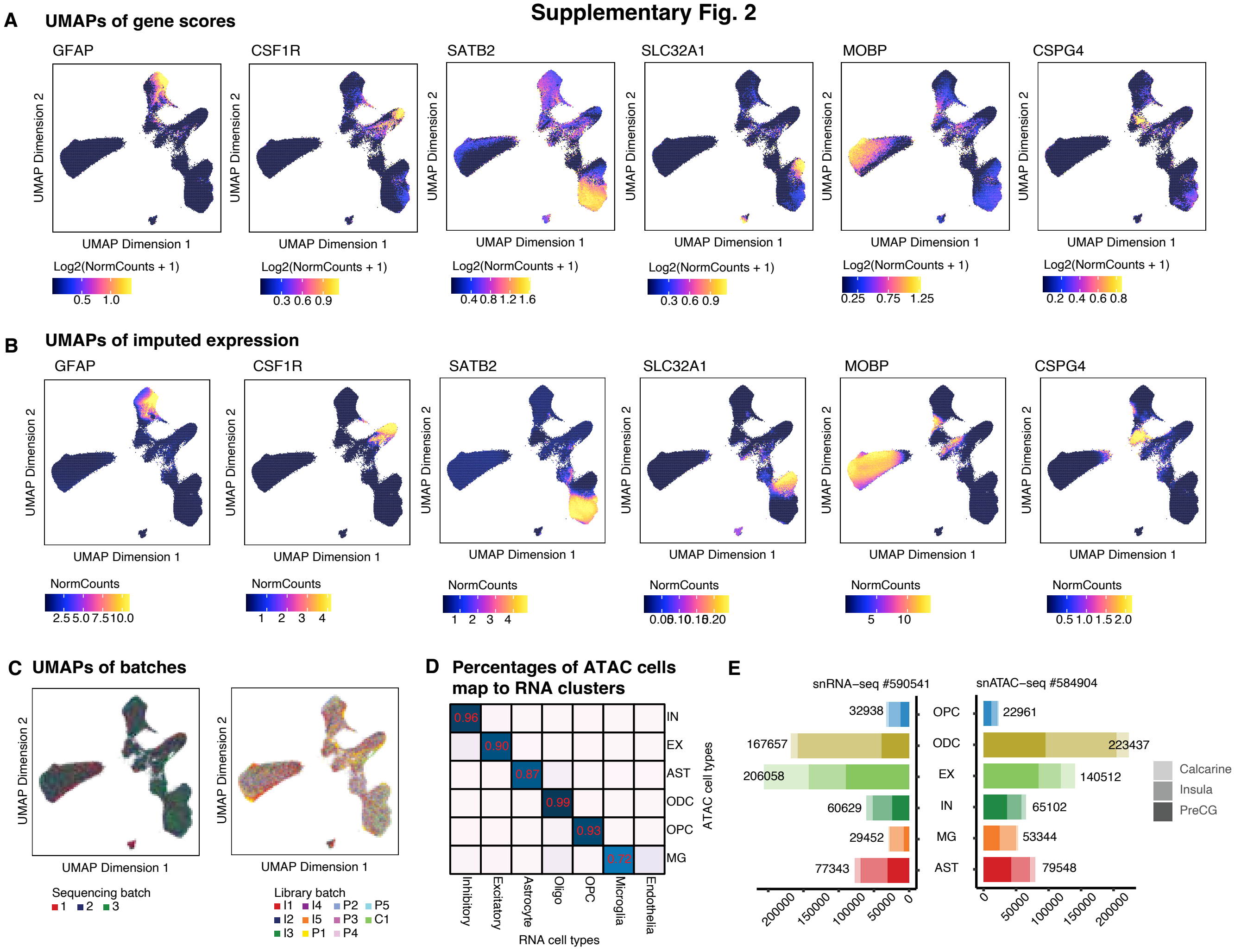

### Supplementary Fig. 3

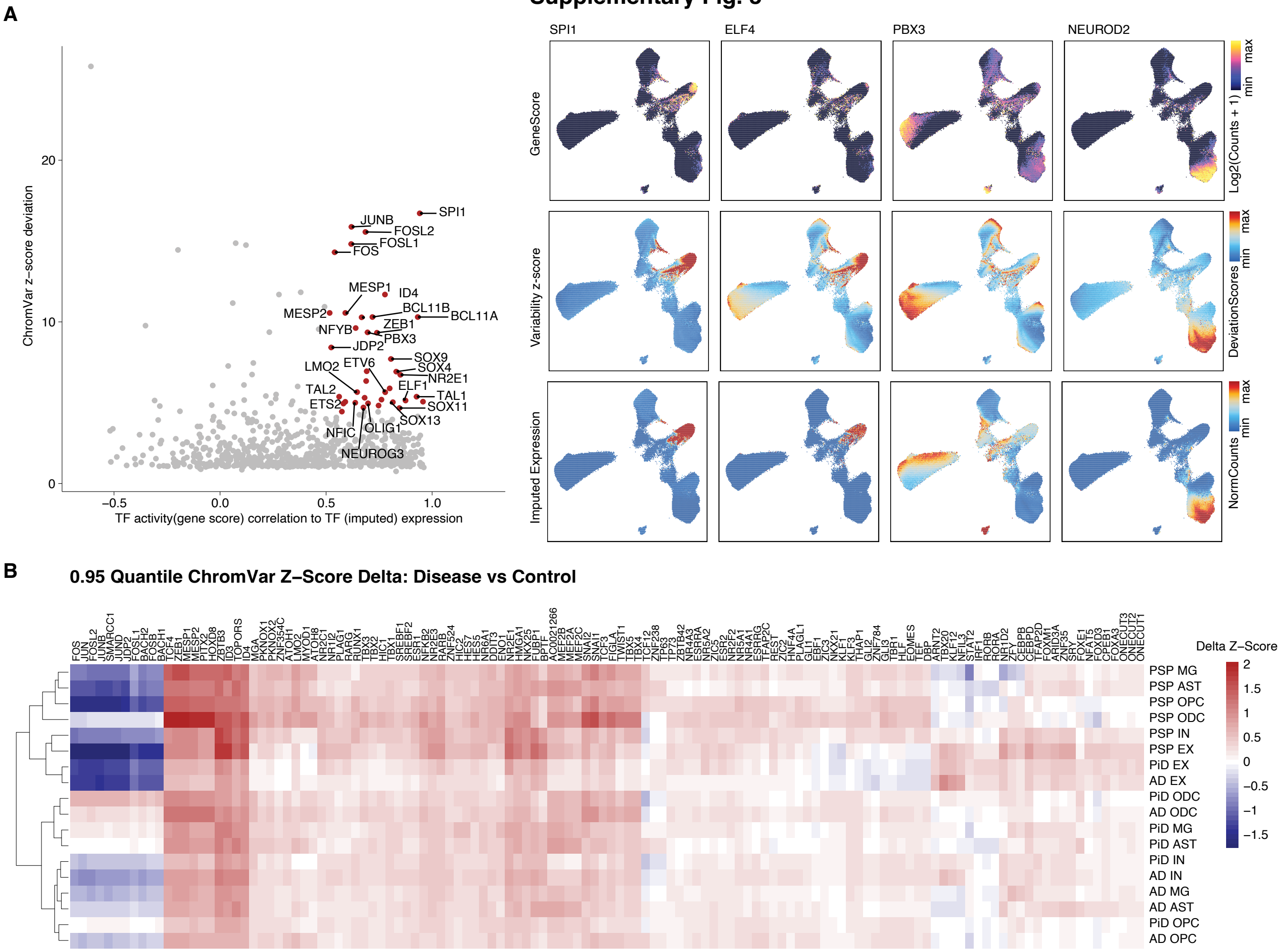

### Supplementary Fig. 4

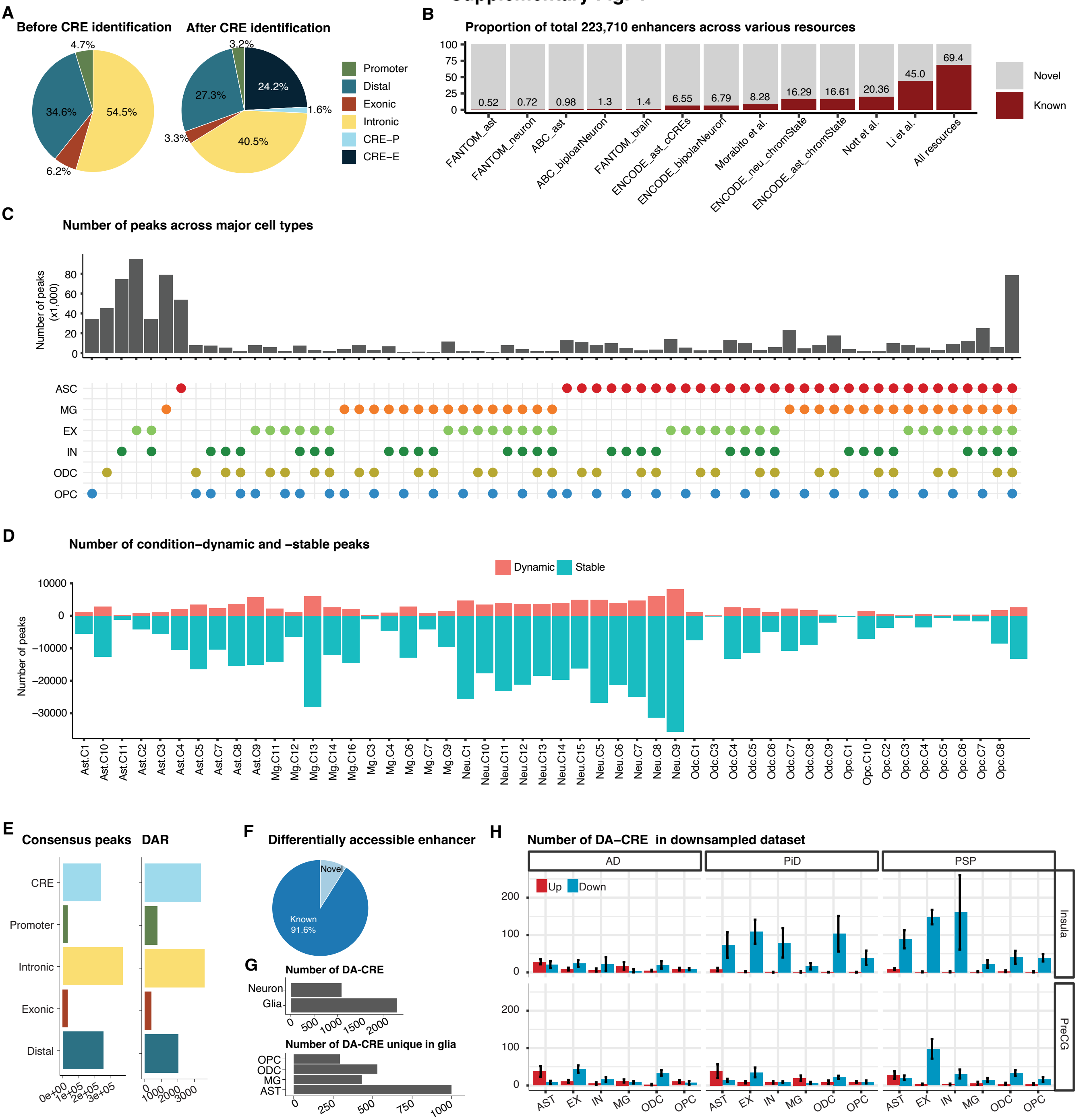

### Supplementary Fig. 5

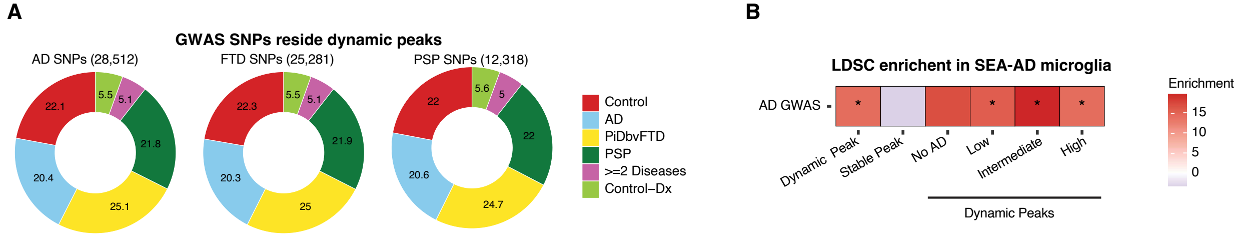

### Supplementary Fig. 6

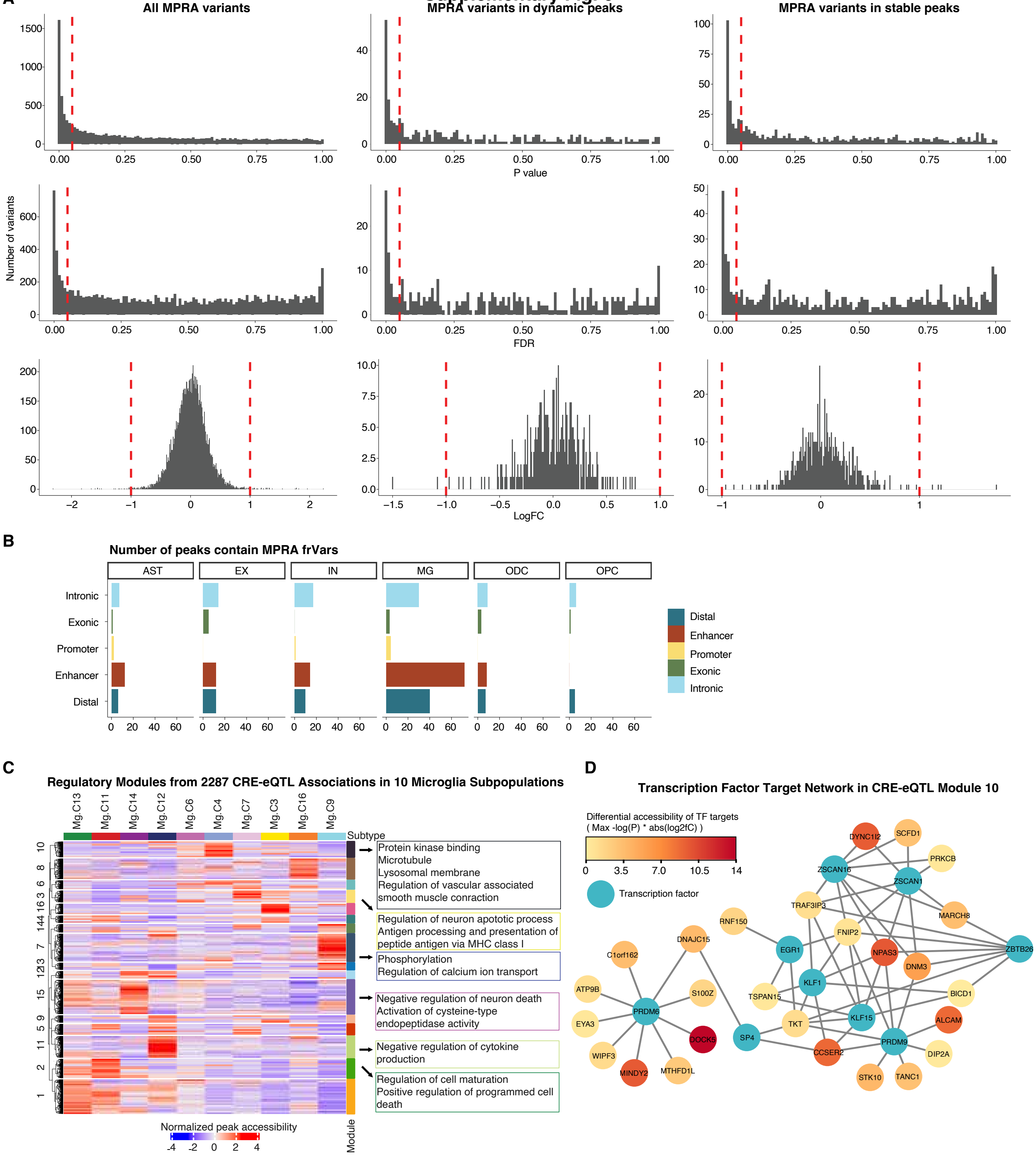

**Supplementary Fig. 7**

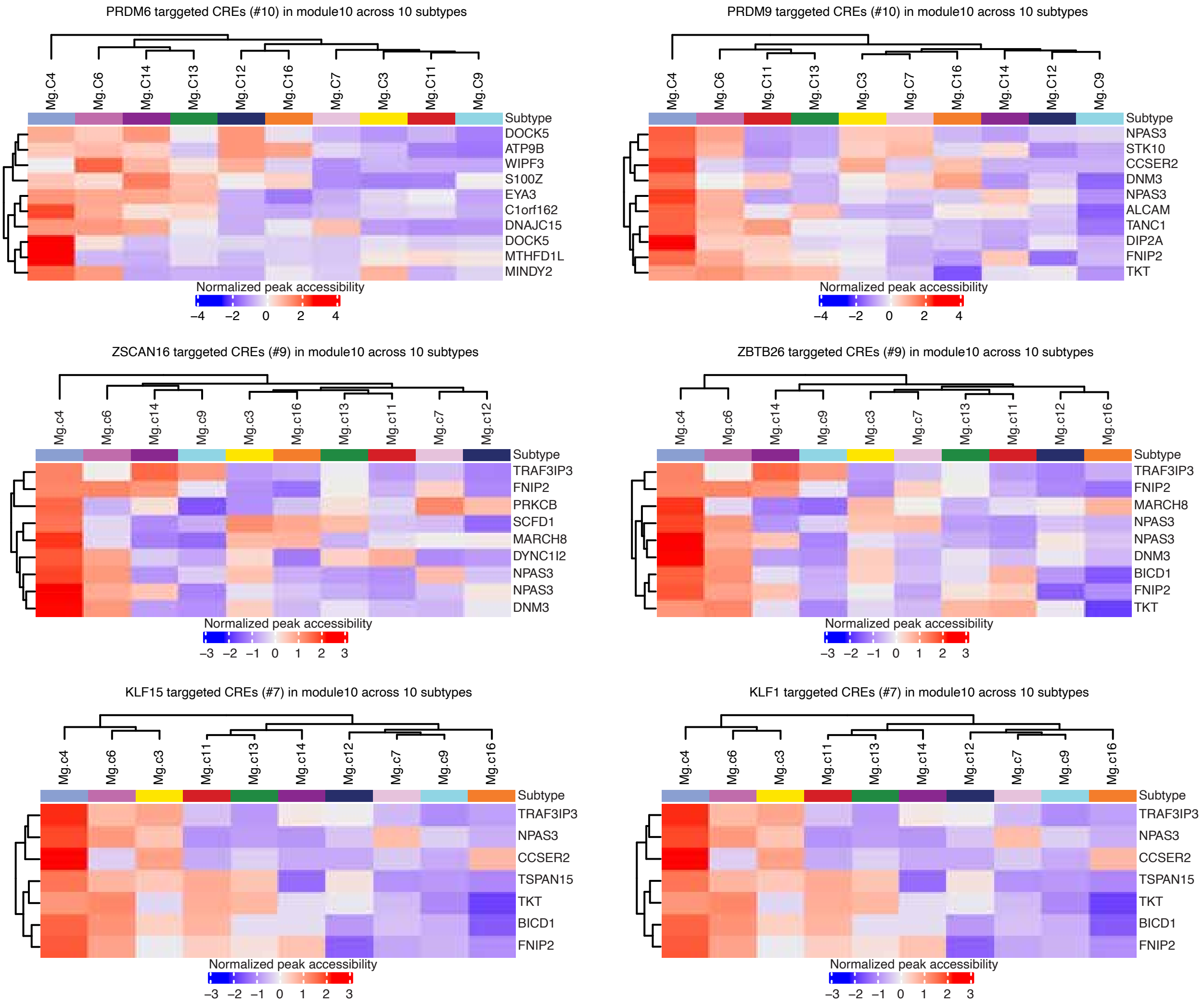

**Supplementary Fig. 8**

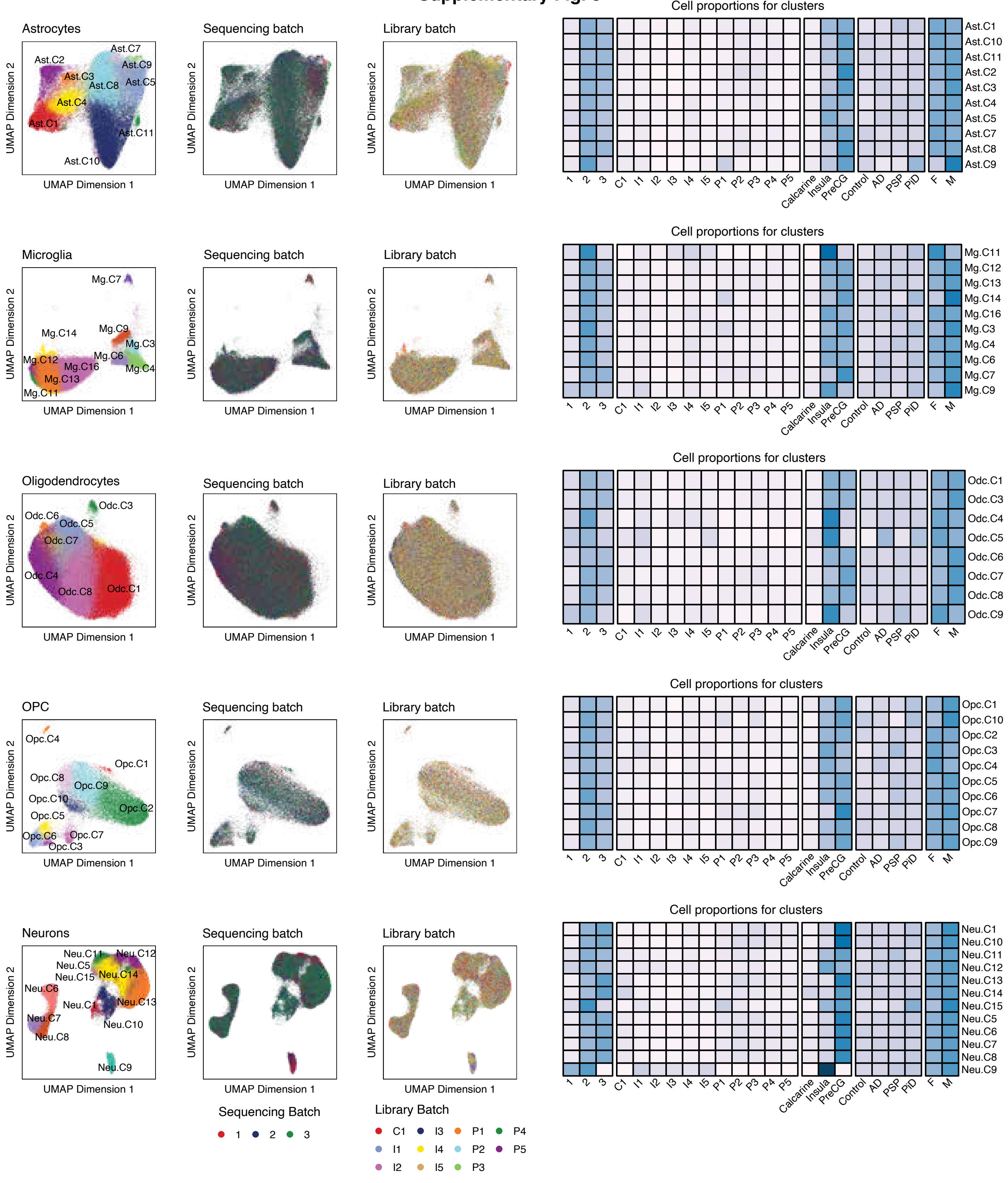

### Supplementary Fig. 9

**A** Cell percentage within cell type and across all cells

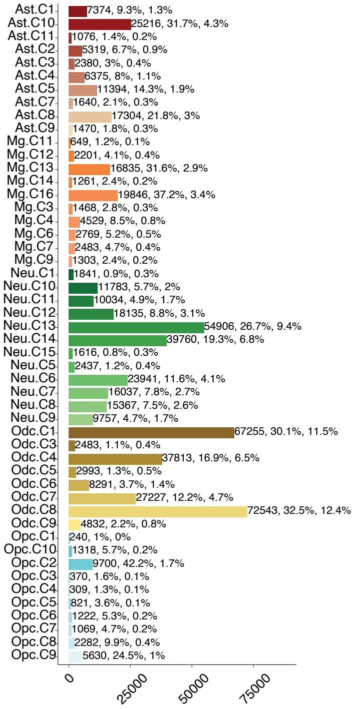


**B** Marker gene scores identified across subclusters

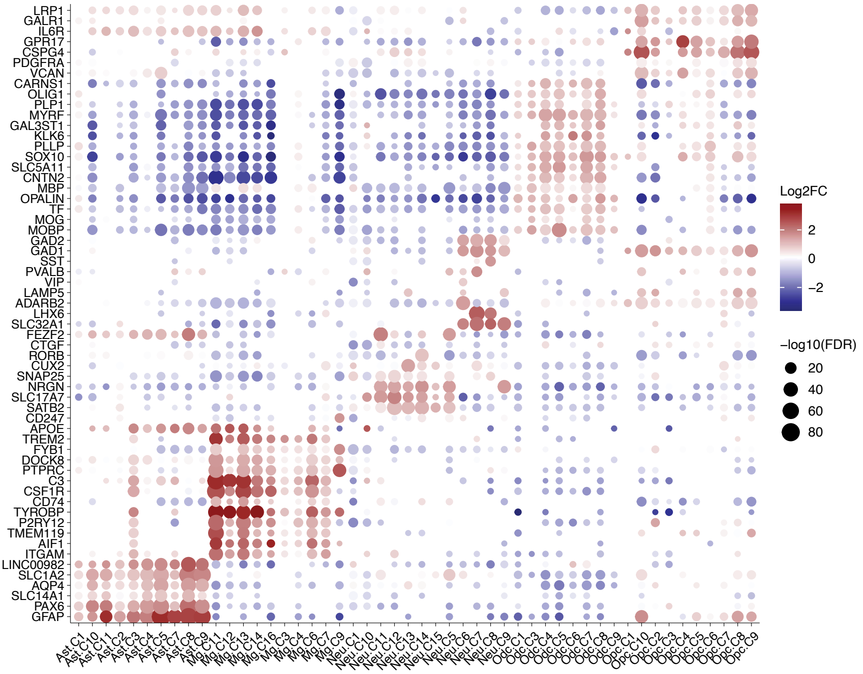


**C** SOX10 gene score

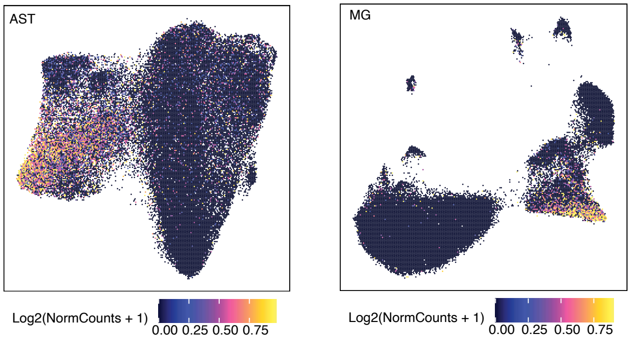

### Supplementary Fig. 10

**A** PSP, DAPI, GFAP-protein, PLP1, SOX10

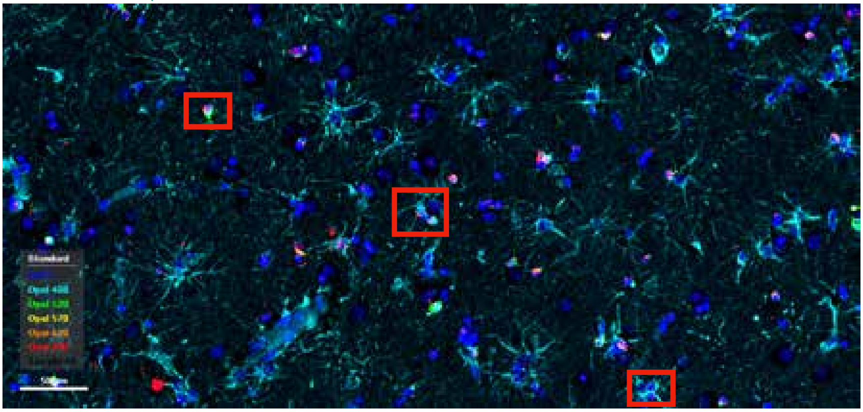

GFAP, SOX10 GFAP, SOX10 PLP1, SOX10

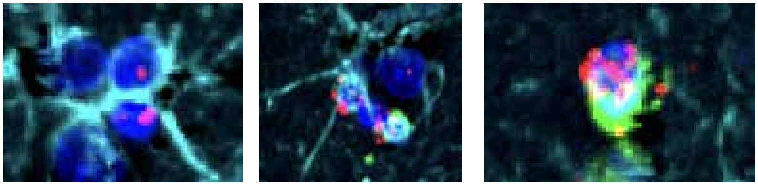

**B** Control, DAPI, GFAP-protein, PLP1, SOX10

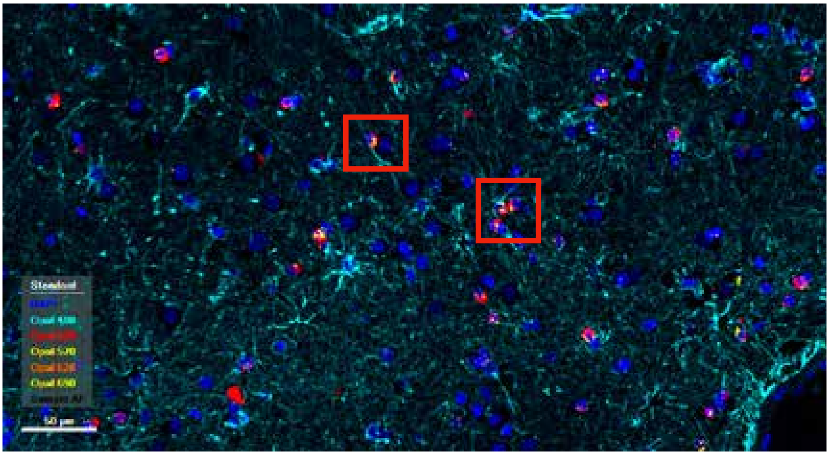

GFAP, PLP1, SOX10

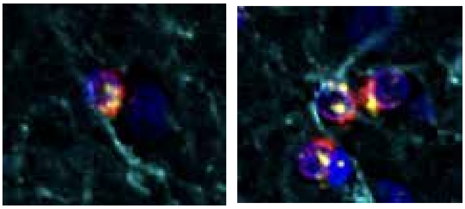

### Supplementary Fig. 11

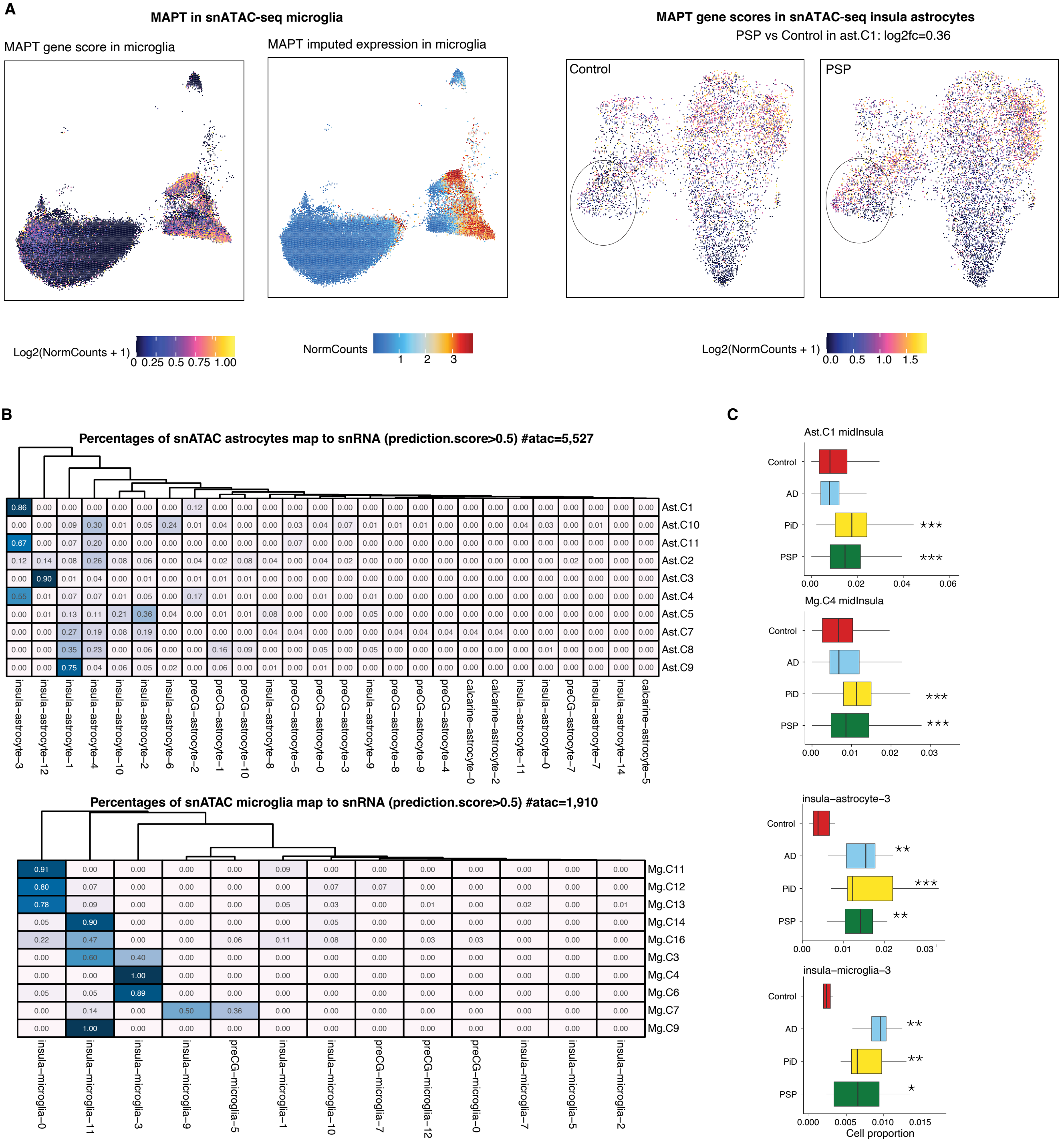

### Supplementary Fig. 12

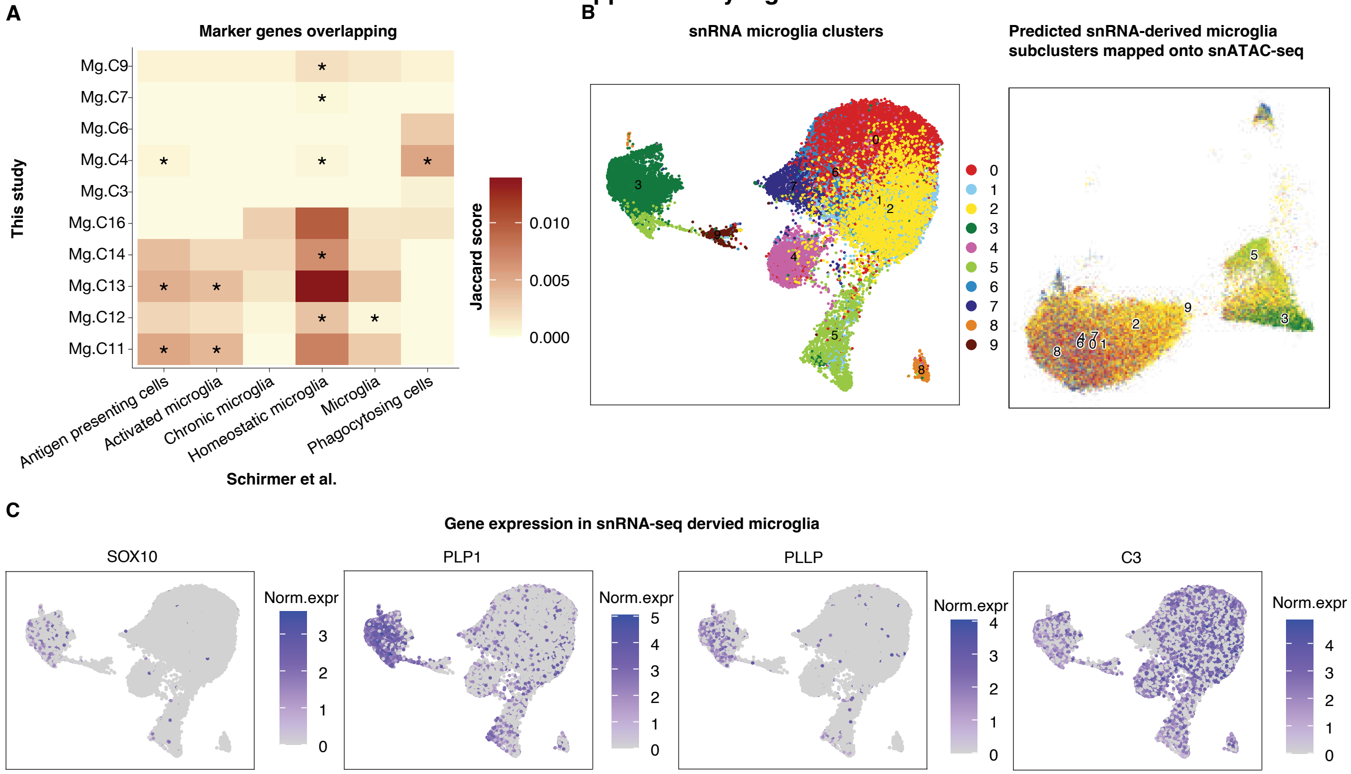

### Supplementary Fig. 13

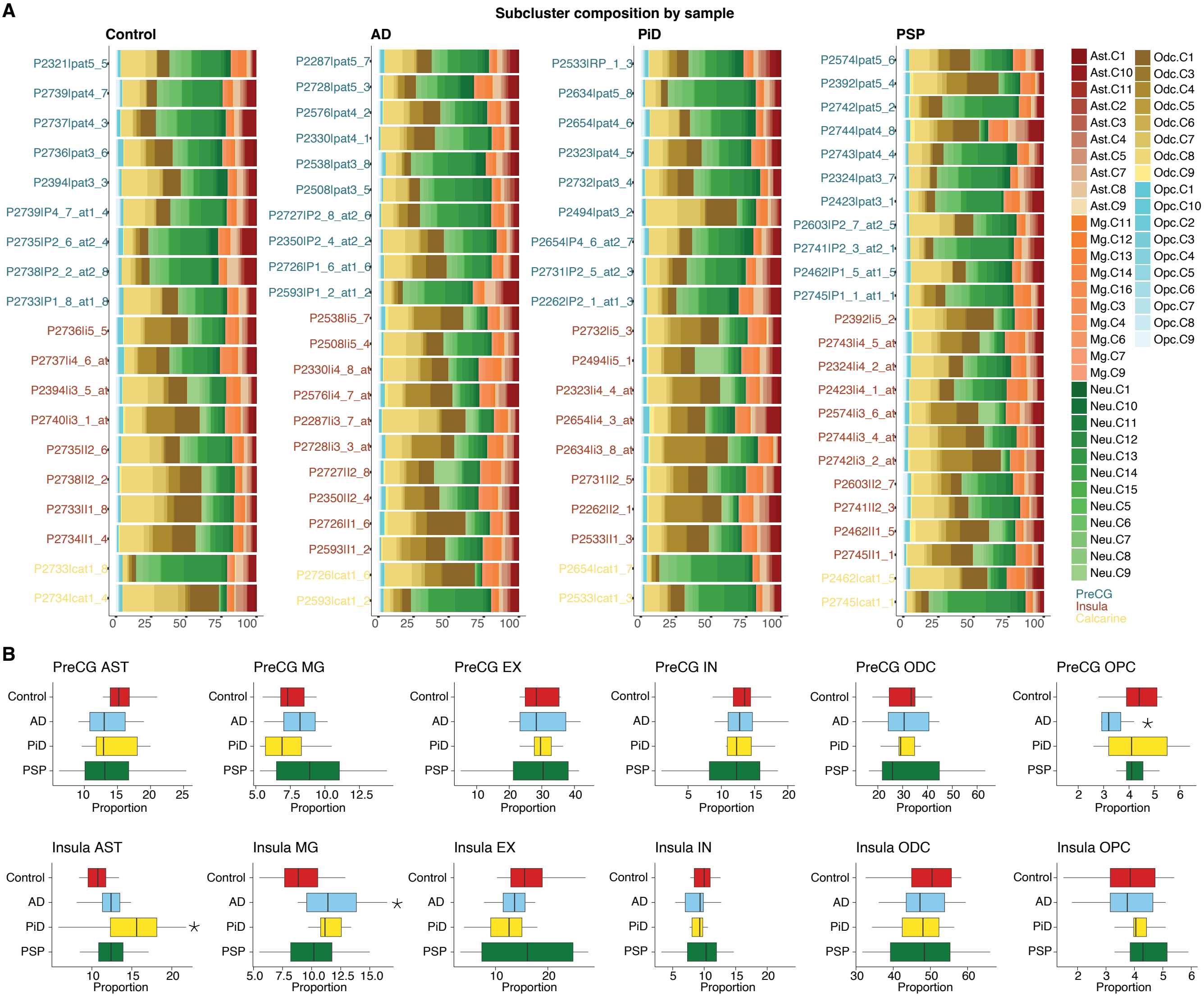

### Supplementary Fig. 14

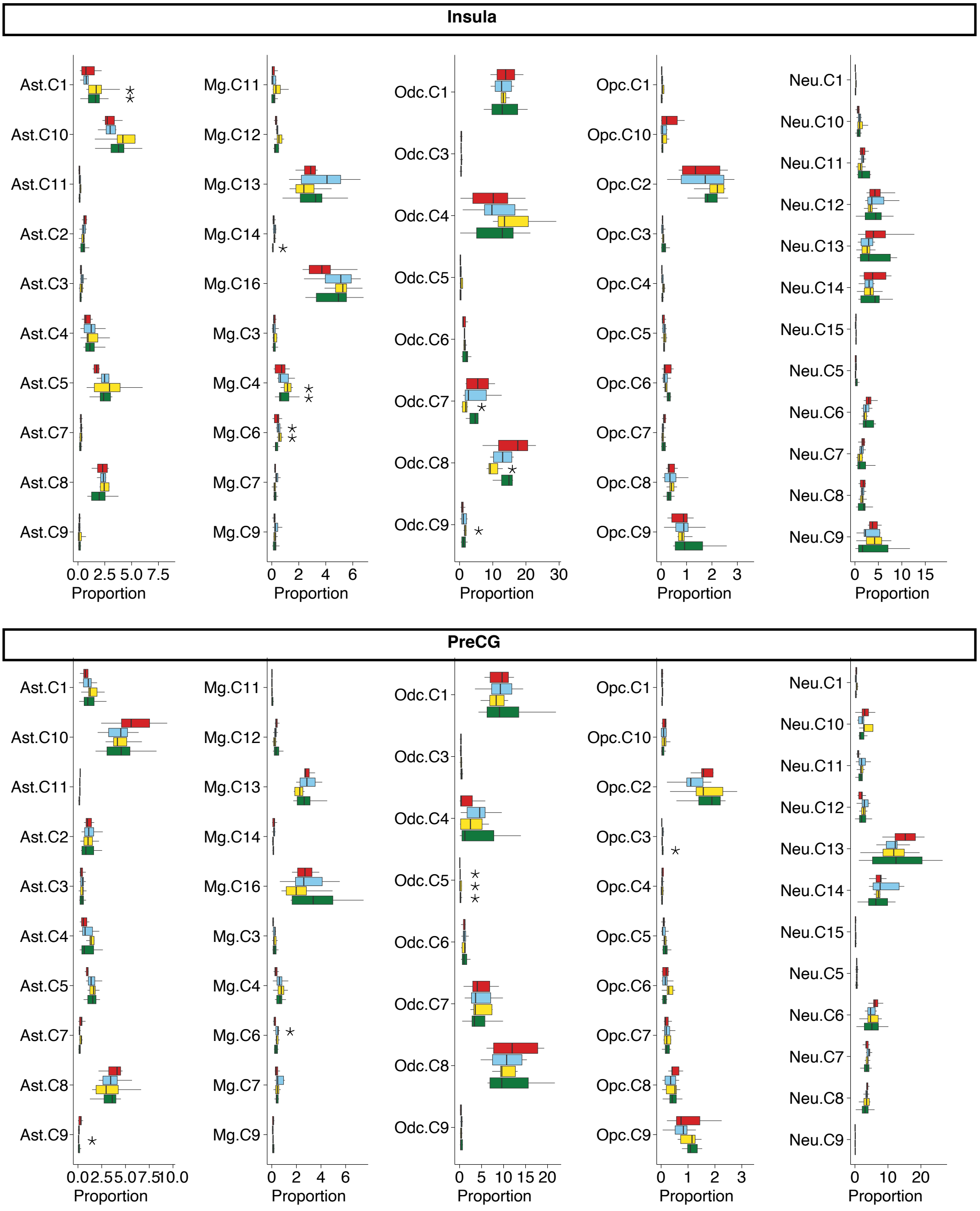

### Supplementary Fig. 15

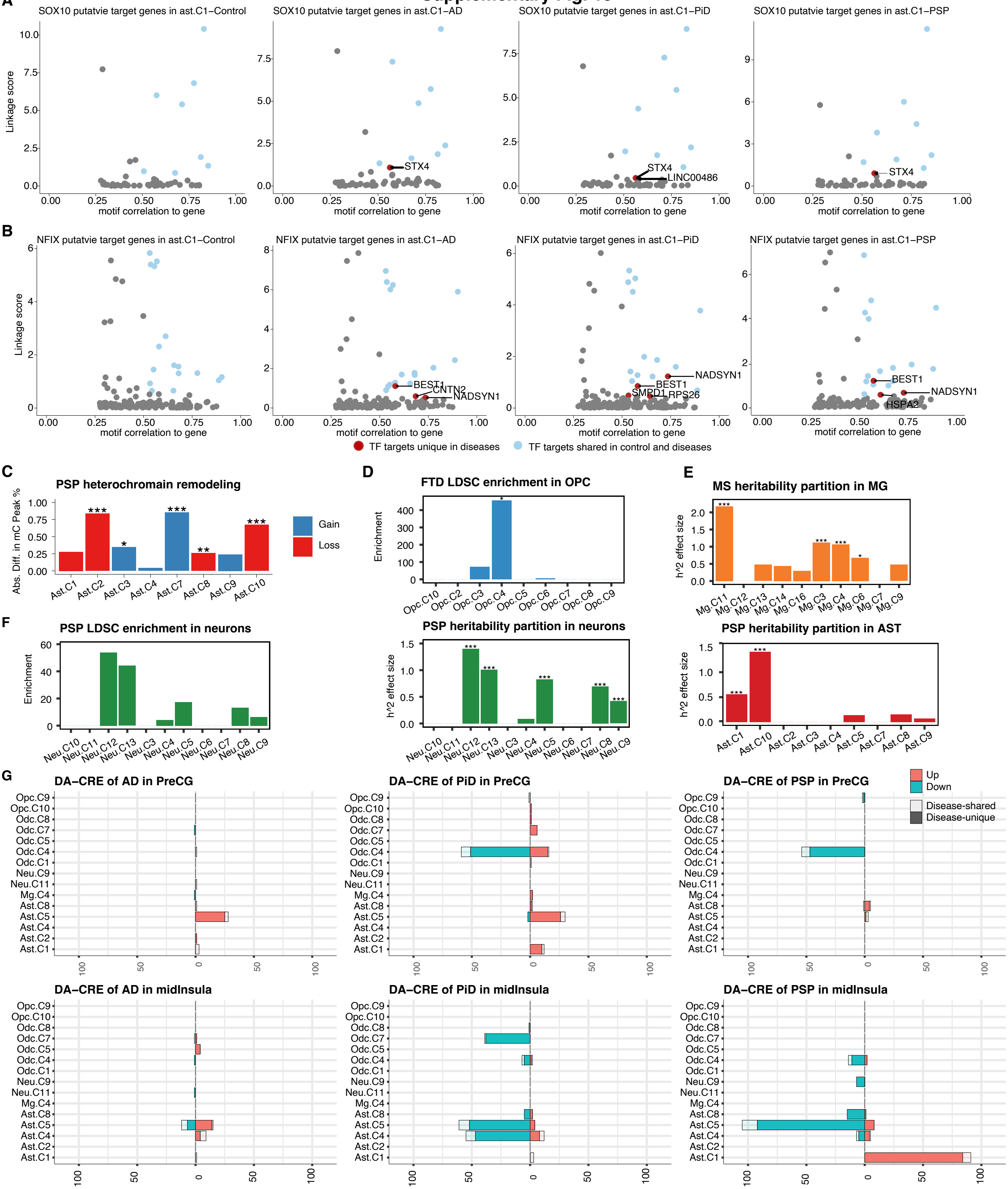

### Supplementary Fig. 16

**A**

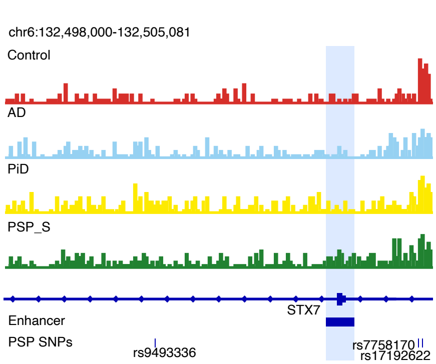


**B**

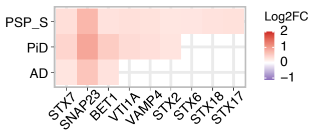
